# Liquid-liquid phase separation and liquid-to-solid transition mediate α-synuclein amyloid fibril containing hydrogel formation

**DOI:** 10.1101/619858

**Authors:** Soumik Ray, Nitu Singh, Satyaprakash Pandey, Rakesh Kumar, Laxmikant Gadhe, Debalina Datta, Komal Patel, Jaladhar Mahato, Ambuja Navalkar, Rajlaxmi Panigrahi, Debdeep Chatterjee, Siddhartha Maiti, Sandhya Bhatia, Surabhi Mehra, Ajay Singh, Juan Gerez, Arindam Chowdhury, Ashutosh Kumar, Ranjith Padinhateeri, Roland Riek, G Krishnamoorthy, Samir K Maji

## Abstract

α-Synuclein (α-Syn) aggregation and amyloid formation is directly linked with Parkinson’s disease (PD) pathogenesis. However, the early events involved in this process remain unclear. Here, using *in vitro* reconstitution and cellular model, we show that liquid-liquid phase separation (LLPS) of α-Syn precedes its aggregation. In particular, *in vitro* generated α-Syn liquid-like droplets eventually undergo a liquid-to-solid transition and form amyloid-hydrogel containing oligomers and fibrillar species. Factors known to aggravate α-Syn aggregation such as low pH, phosphomimic substitution, and familial PD mutation also promote α-Syn LLPS and its subsequent maturation. We further demonstrate α-Syn liquid droplet formation *in cells*, under oxidative stress. These cellular α-Syn droplets eventually transform into perinuclear aggresomes, the process regulated by microtubules. The present work provides detailed insights into the phase separation behavior of natively unstructured α-Syn and its conversion to a disease-associated aggregated state, which is highly relevant in PD pathogenesis.

## INTRODUCTION

Liquid-liquid phase separation (LLPS) of biological polymers (protein, RNA) into liquid condensates has emerged as a critical phenomenon in the formation of intracellular “membrane-less” organelles (Alberti, 2017; Banani et al., 2017; Boeynaems et al., 2018; Brangwynne et al., 2009; Hyman et al., 2014; Shin and Brangwynne, 2017). Such examples include nucleoli (Feric et al., 2016), Cajal bodies (Kaiser et al., 2008), PML bodies (Banani et al., 2016) in the nucleus as well as stress granules in the cytoplasm (Lin et al., 2015; Molliex et al., 2015; Riback et al., 2017). These liquid condensates concentrate biomolecules such as proteins and nucleic acids at distinct cellular sites and perform various cellular functions (Shin and Brangwynne, 2017). Due to the lack of a physical barrier, such as membrane, the liquid condensates are able to exchange their components rapidly with the surrounding medium (Hyman et al., 2014; Mitrea and Kriwacki, 2016). Most of the liquid condensates possess common characteristics, which include their formation mechanism as well as their physical properties. For instance, multivalent proteins or nucleic acids associate through weak intermolecular interactions and reach a solubility limit to form liquid condensates (Banani et al., 2017; Shin and Brangwynne, 2017). These condensates are highly mobile, spherical, but get deformed on physical contact, fuse and eventually relax back to their spherical shape (Brangwynne et al., 2009; Brangwynne et al., 2011; Molliex et al., 2015; Nott et al., 2015). Several proteins undergoing LLPS, however, contain intrinsically disordered regions (IDRs) that are closely associated with prion-like domains (PLDs) and low complexity domains (LCDs) (Hughes et al., 2018; Maharana et al., 2018; Pak et al., 2016; Wang et al., 2018), where the amino acid variance is extremely low (Burke et al., 2015; Elbaum-Garfinkle et al., 2015; Zhang et al., 2015). These IDRs drive LLPS by weak, multivalent interactions between the protein molecules, thus allowing various homotypic and heterotypic interactions of proteins and other biomolecules (Li et al., 2012; Lin et al., 2015; Mitrea et al., 2016; Molliex et al., 2015; Nott et al., 2015).

Many proteins which initially form highly mobile liquid condensates become more viscoelastic and rigid during time and eventually form a gel-like state that is unable to exchange its component molecules with the surrounding (Molliex et al., 2015; Murakami et al., 2015; Patel et al., 2015; Wegmann et al., 2018). This transition could either be due to entanglement of biopolymers or stronger association of proteins leading to fibril formation as reported for many protein condensates associated with neurological disorders such as FUS, TDP-43, Tau and hnRNPA1 (Ambadipudi et al., 2017; Conicella et al., 2016; Molliex et al., 2015; Patel et al., 2015; Wegmann et al., 2018). It has been suggested that in these cases, phase separation might increase the nucleation rate for protein aggregation into amyloid fibrils (Lin et al., 2015; Pak et al., 2016; Xiang et al., 2015). Following this concept, in the study presented, we hypothesized that α-Syn aggregation associated with PD pathogenesis (Spillantini et al., 1997; Volles and Lansbury, 2003) might involve LLPS.

α-Synuclein (α-Syn) is a natively unstructured protein (Eliezer et al., 2001), and its aggregation into cytotoxic oligomers and amyloid fibrils are associated with PD pathogenesis (Goedert, 2001). α-Syn aggregation involves nucleation-dependent polymerization (Conway et al., 2000), where the unstructured protein first forms partially folded intermediates (Uversky et al., 2001) followed by oligomerization and fibrillation into amyloids. Moreover, the mutations of α-Syn associated with early onset aggressive form of familial PD (Kruger et al., 1998; Pasanen et al., 2014; Zarranz et al., 2004), are known to modulate its aggregation (Conway et al., 1998; Ghosh et al., 2014; Greenbaum et al., 2005), further supporting its role in PD pathogenesis. Although the mechanism of α-Syn aggregation is an area of extensive research, most studies are focused on delineating the end-stage of the aggregation process; however, the early events (involving the formation of transient species) leading to the formation of aggregates is not well established. The primary structure of α-Syn contains three distinct domains: the N terminal region, an aggregation-prone “non-amyloid component (NAC)” domain and a flexible C-terminal domain. Although the NAC region primarily drives α-Syn aggregation (Giasson et al., 2001), the majority of familial mutations are, however, at the N-terminal region highlighting its importance in α-Syn misfolding and aggregation as demonstrated by the cryo-electron microscope structure of α-Syn fibrils (Guerrero-Ferreira et al., 2018; Li et al., 2018a). Further, the N-terminal domain possesses two LCDs suggesting that α-Syn might undergo phase separation under appropriate conditions.

We demonstrate here LLPS and liquid droplet formation by α-Syn at a high local concentration (in the presence of molecular crowder), which is greatly promoted by various known conditions associated with PD pathogenesis. Detailed biophysical and structural characterization revealed that the α-Syn droplets undergo a liquid to solid-like transition, eventually transforming into a gel state containing fibrillar aggregates and oligomers. Structural studies establish that low-complexity region of N-terminus and the central hydrophobic NAC domain majorly drive α-Syn LLPS. In cells, α-Syn formed liquid droplets under oxidative stress, which subsequently mature and transform into solid-like aggregates forming aggresomes, the process regulated by microtubules. Our findings establish that α-Syn phase separates into liquid droplets, which act as an initial step towards α-Syn aggregation associated with PD pathology.

## RESULTS

### Liquid-liquid phase separation of α-Syn *in vitro*

Proteins with intrinsically disorder regions (IDRs) and low complexity domains (LCDs) are known to phase separate *in vitro* when the local concentration of the protein molecules becomes high in the presence of a molecular crowding agent (Fonin et al., 2018). We used bioinformatics tool such as SMART (Letunic and Bork, 2018) and IUPred2 (Meszaros et al., 2018) algorithms to examine the presence of IDRs and LCDs, respectively, in α-Syn. As indicated in Figure 1A, SMART analyses predicted two LCDs (residues 10-23 and 63-78) in the primary sequence of α-Syn (upper panel), and IUPred2 revealed disorderness in the segment of 26-28, 54-56 and 99-140 (lower panel). We, therefore, hypothesized that α-Syn might undergo LLPS under appropriate conditions. We examined the phase separating behavior of α-Syn at different protein concentrations in 20 mM phosphate buffer, pH 7.4, in the presence and absence of 10% polyethylene glycol (PEG)-8000 as a molecular crowding agent. Incubation of α-Syn at 37°C in the presence of PEG showed the formation of liquid-like droplets at high protein concentrations (≥200 μM) (Figure 1B). Increasing the molecular crowding further decreased the critical concentration of α-Syn to phase separate. For example, in the presence of 15 – 20% PEG even at 10 μM protein concentration liquid-like droplets are formed (Figures 1D and S1A), while the control with PEG alone in phosphate buffer, pH 7.4 did not show any droplet formation (data not shown). This suggests that an increased local concentration of α-Syn by crowding is sufficient to initiate α-Syn LLPS under physiological condition. α-Syn LLPS was further confirmed using light scattering, which showed an increase in scattering after 48 h, as anticipated for phase-separated samples (Figure S1B). Fluorescence imaging studies with orthogonal-plane sectioning using fluorescein isothiocyanate (FITC)-labeled α-Syn confirmed that these liquid droplets indeed contain α-Syn molecules (Figures 1C and S1C).

**Figure 1:**
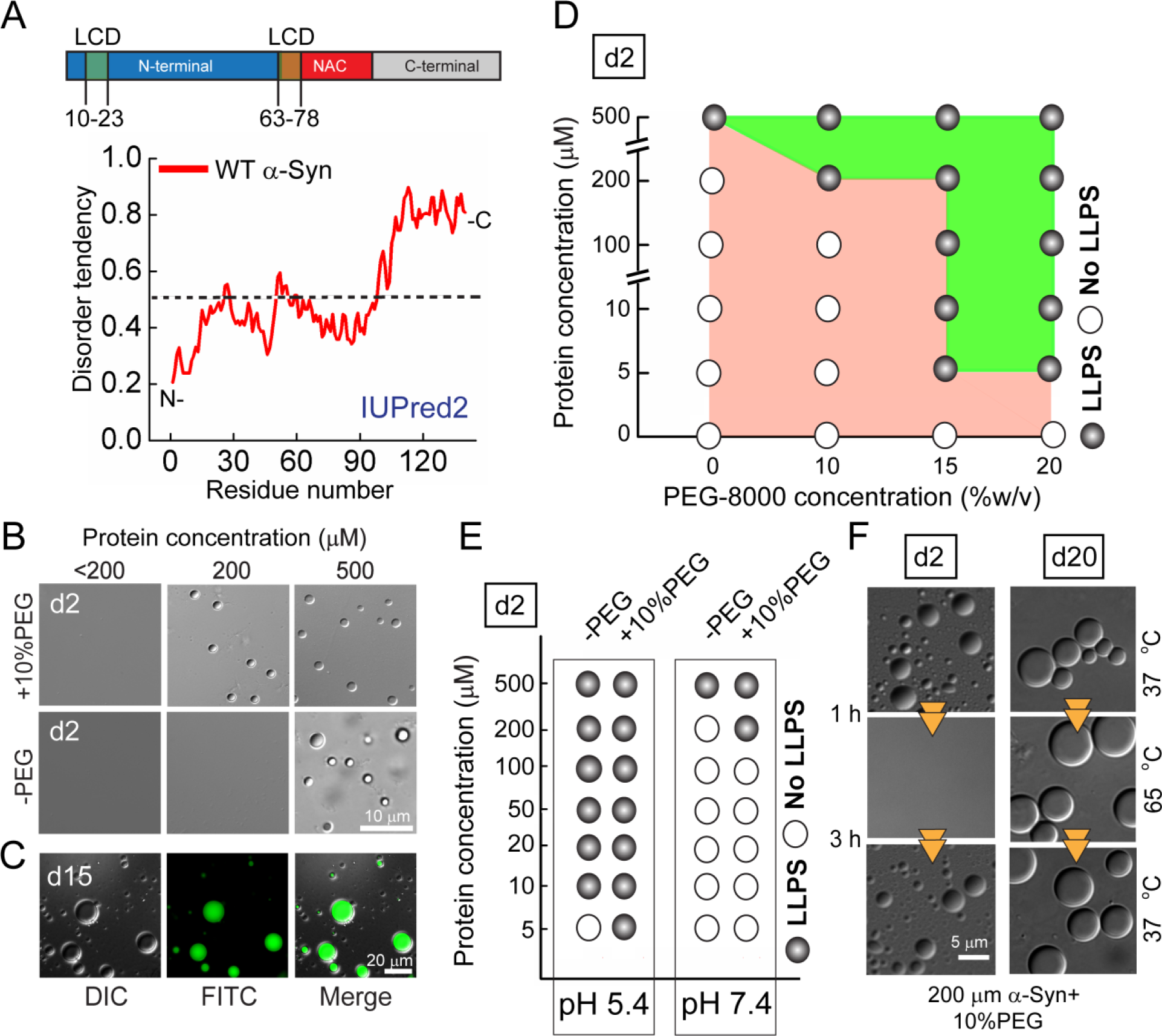
α-Syn undergoes liquid-liquid phase separation *in vitro*. **A**. *In silico* analysis of the primary sequence of WT α-Syn using SMART and IUPred2. LCDs were predicted by SMART (top panel) and the disorder tendency was analyzed using IUPred2 prediction (bottom panel). **B.** DIC images of α-Syn phase separated droplets at different protein concentrations in the presence and absence of molecular crowder, PEG-8000. **C.** Fluorescence images of FITC-labeled α-Syn (200 μM) phase separated droplets formed in presence of 10% PEG-8000 at day 15 (d15). **D**. Regime diagram illustrating phase separation of α-Syn at different protein and PEG-8000 concentrations. **E**. Regime diagram of α-Syn at pH 5.4 and 7.4. Filled circles (●) indicate phase separation and open circles (○) as no phase separation. **F.** DIC images of α-Syn (200 μM) in presence of 10 % PEG, at day 2 (d2) and day 20 (d20), demonstrating the reversible and irreversible nature of the droplets, respectively upon heating (65 °C) and cooling (37°C).

In order to explore the phase space of the LLPS, we compared phase separation of α-Syn at various concentrations and at two different pHs, 5.4 and 7.4 (in the absence and presence of 10 % PEG). At pH 5.4, α-Syn formed droplets even at concentrations of ≤10 μM in the absence of PEG, and at 5 μM in the presence of 10% PEG (Figures 1E and S1D). As many protein droplets showed dependency on temperature (for the formation and reversibility) (Falahati and Wieschaus, 2017; Nott et al., 2015; Reichheld et al., 2017; Schuster et al., 2018), we also tested temperature-dependent α-Syn phase separation (i.e. 4 ºC, 18 ºC, 25 ºC and 37 ºC) using 200 μM protein concentration. α-Syn liquid droplets were observed at all temperatures (except at 4 °C) after 48 h of incubation (Figure S1E). To examine the temperature reversibility, we subjected the freshly formed α-Syn liquid droplets to increasing temperature and observed that the newly formed droplets disappeared when the temperature was raised to 65 °C, while they reappeared within 1 h upon cooling to 37 °C (Figures 1F and S1F). The disappearance of liquid droplets upon heating and reappearance upon cooling highlights their reversible, liquid-like nature. However, 20 days aged droplets incubated at 37 ºC did not show this temperature reversibility suggesting the appearance of a rigid-like hydrogel state (Figures 1F and S1F). This shows a time-dependent change of the physical properties of phase-separated α-Syn droplets from a liquid-liquid to a solid-like state. Similar characteristics have also been observed for liquid droplets formed by FUS (Patel et al., 2015), TDP-43 (Conicella et al., 2016) and Tau (Wegmann et al., 2018) undergoing a transition from the reversible liquid state to an aberrant gel-like state of irreversible aggregates.

### Dynamics of α-Syn molecules inside the phase separated liquid droplets

To understand the dynamics of α-Syn molecules, the droplet formation and its maturation was monitored over time (20 days). The time-dependent analysis of α-Syn LLPS under the microscope showed that during maturation, the size of the droplets increased (Figure 2A and Video S1), which could be due to Ostwald ripening or droplet fusion (Berry et al., 2015; Voorhees, 1992). Representative time-lapse images show coalescence of two droplets forming a larger sized droplet (Figure 2B) further supporting the existence of droplet fusion. Subsequently, to characterize the dynamic nature of the droplets, we performed fluorescence recovery after photobleaching (FRAP) experiments using droplets containing 10 % rhodamine-labeled α-Syn. FRAP studies revealed that fluorescence recovery of the bleached region of α-Syn droplets is very rapid at the beginning of their formation (i.e. day 2-old droplets show a 96 % recovery and a diffusion coefficient of 0.584 μm^2^s^−1^), which significantly decreased after 20 days of aging, where only ~7.5 % recovery was observed (Figures 2C and D). The average diffusion coefficients calculated for day 5 and day 10 droplets were 0.2402 and 0.187 μm^2^s^−1^, respectively. The diffusion coefficients for day 15 and 20 could not be calculated as the % recovery showed no exponential increase with time. The progressive rigidity (low recovery from FRAP) is attributed to increased viscoelasticity and decreased molecular diffusion of the protein molecules inside the droplets during their maturation (Kato et al., 2012). Furthermore, to confirm the liquid to solid-like transition of the α-Syn droplets, we measured time-resolved fluorescence anisotropy decay from inside and outside of the FITC-tagged α-Syn phase separated droplets. Fluorescence anisotropy decay is expected to be very fast when α-Syn is in the liquid state but is slow when in the solid-like state (Sahay et al., 2016). P1 (inside the droplet) and P2 (outside the droplet) are the reference pixels under the microscopic field from which the fluorescence anisotropy decay was obtained (Figure 2E, left panel). The fluorescence anisotropy decay inside the droplet was slower compared to outside, which clearly reflects the rigidity of the protein molecules within the droplet compared to the outside (bulk phase). The α-Syn molecules inside the day10 old droplets, however, were more rigid compared to day 2 droplets (freshly formed), as seen by the decrease in the fluorescence anisotropy decay (Figure 2E, right panel). These observations are in accordance with FRAP recovery data, which asserts the transition from liquid to solid-like state during incubation. It should be noted here that the FRAP data reports on the translational diffusion of the fluorophore, however, the time-resolved fluorescence anisotropy provides information on the rotational motions (Sahay et al., 2016). The observed results, therefore, clearly suggest that both the translational and rotational motions of the fluorophore are slowed as the droplets mature over time.

**Figure 2:**
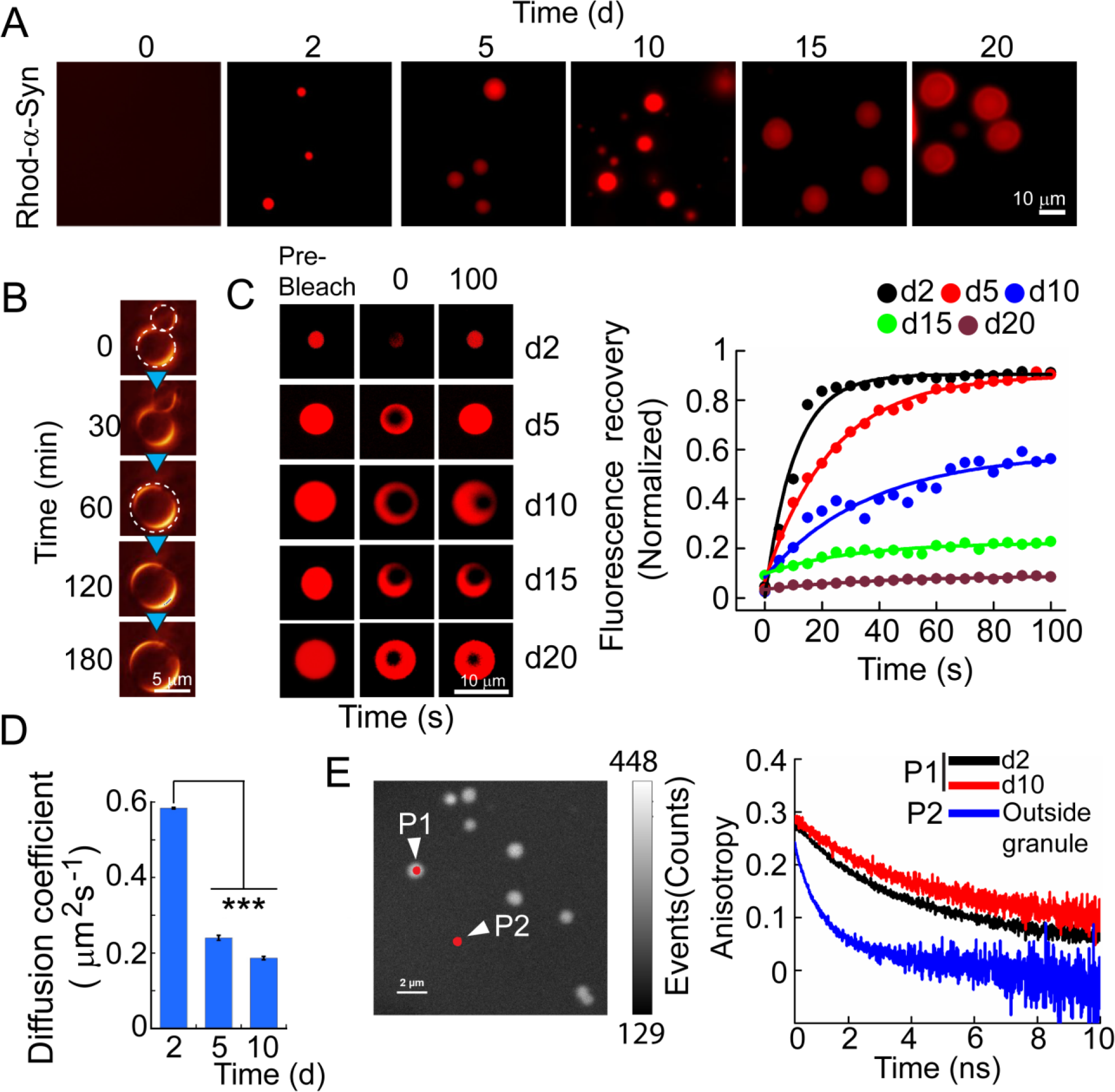
Dynamics of α-Syn in LLPS slows down with time. **A.** Fluorescence images showing the maturation of NHS-rhodamine labeled α-Syn droplets over time. **B.** Time-lapse images of α-Syn droplet showing the fusion of two droplets and formation of a larger single droplet over the time, represented in pseudocolor. **C.** FRAP measurements of α-Syn droplets at different times to measure the change in dynamics of droplets at day 2 (d2), day 5 (d5), day 10 (d10), day 15 (d15) and day 20 (d20). A representative droplet at indicated time points and corresponding FRAP intensity curves are shown alongside. **D.** Diffusion coefficients at indicated time points are shown. Notably, the diffusion coefficient could not be calculated for day 15 and day 20 droplets due to negligible recovery after photobleaching. Area of ROI, ω^2^ = 9 μm^2^; n=3, values represent mean ± s.e.m (***p ≤ 0.001). **E**. Left panel: Representative image of FITC-labeled α-Syn droplets analyzed with time-resolved fluorescence anisotropy decay. P1 and P2 are points from inside and outside of the droplets, respectively, used for the analysis. Right panel: Fluorescence anisotropy decay curves demonstrating a delayed decay for day 10 (d10) droplets compared to day 2 (d2), indicating increased rigidity of α-Syn molecules during droplet maturation. All the experiments were carried out with 200 μM of the protein in presence of 10 % PEG.

### Parkinson’s disease-promoting factors accelerate α-Syn aggregation and droplet formation

PD is mostly a sporadic disorder where environmental and cellular factors play major roles in the disease pathogenesis (Cookson, 2005). In this context, factors including metal ions, interaction with a lipid membrane, phosphorylation of Ser129, and familial mutations are known to play a critical role in aggregation of α-Syn in the diseased brain as well as *in vitro* (Breydo et al., 2012). Similarly, it has been shown that factors that modulate the assembly of protein or solubility, such as post-translational modification and pH play a critical role in modulating LLPS (Ambadipudi et al., 2017; Monahan et al., 2017; Nott et al., 2015; Riback et al., 2017). In line with this correlation is the finding that lowering the pH promotes both α-Syn LLPS (Figures 1E and S1D) and α-Syn aggregation (Buell et al., 2014). To further strengthen this apparent correlation, we asked whether these PD-associated factors also affect the phase separation behavior of α-Syn. We first tested the aggregation kinetics of α-Syn using thioflavin T (ThT) fluorescence assays in the presence and absence of these modulators. Cu^+2^, Fe^+3^, liposomes, and S129E phosphomimetic accelerated α-Syn aggregation compared to α-Syn alone, while the fibril formation interfering dopamine (Conway et al., 2001; Jha et al., 2017) delayed the aggregation kinetics (Figure 3A). As expected the aggregated state showed amyloid-like fibrils under all the conditions as confirmed by TEM (Figure S2A). When we studied the liquid droplet formation in the presence of the modulators (Cu^+2^, Fe^+3^, and liposomes), faster liquid droplet formation (within 24 h) without any requirement of the molecular crowder, PEG was observed (Figures 3B and S2B). Although the phosphomimetic mutant S129E α-Syn could not form droplets in the absence of molecular crowder, it phase separated at a faster rate in the presence 1-10% PEG (24 h) when compared to the WT protein (Figures 2A and 3B). Apart from the faster rate of liquid droplet formation, many of these factors also reduced the critical concentration of α-Syn for LLPS. For example, α-Syn formed liquid droplets at 5 μM protein concentration in the presence of 100 μM Cu^+2^ or 1 mM liposomes (Figure S2B). Also, in the presence of 15 % PEG, the S129E variant formed droplets at concentrations as low as 5 μM. The data, therefore, suggest that these factors lower the critical concentration of α-Syn phase separation due to an increased propensity for intermolecular interaction between α-Syn molecules in these conditions. A53T is a familial mutant of α-Syn associated with early-onset PD (Conway et al., 1998; Polymeropoulos et al., 1997), which is known to aggregate faster *in vitro* than WT (Narhi et al., 1999). Similar to WT α-Syn, A53T mutant also phase separated in the presence of PEG after 48 h (Figure 3B), however, the propensity of droplet formation for A53T was higher compared to WT, as confirmed by the light scattering measurements (Figure S2D). Furthermore, the A53T mutant showed significantly increased number and larger sized droplets compared to WT α-Syn (Figure S2E). In contrast, in presence of dopamine, no phase separation occurred even after extended incubation (7 days) of 200 μM α-Syn in presence of 10% PEG (Figure S2C); thus, providing a direct link between LLPS and aggregation.

**Figure 3:**
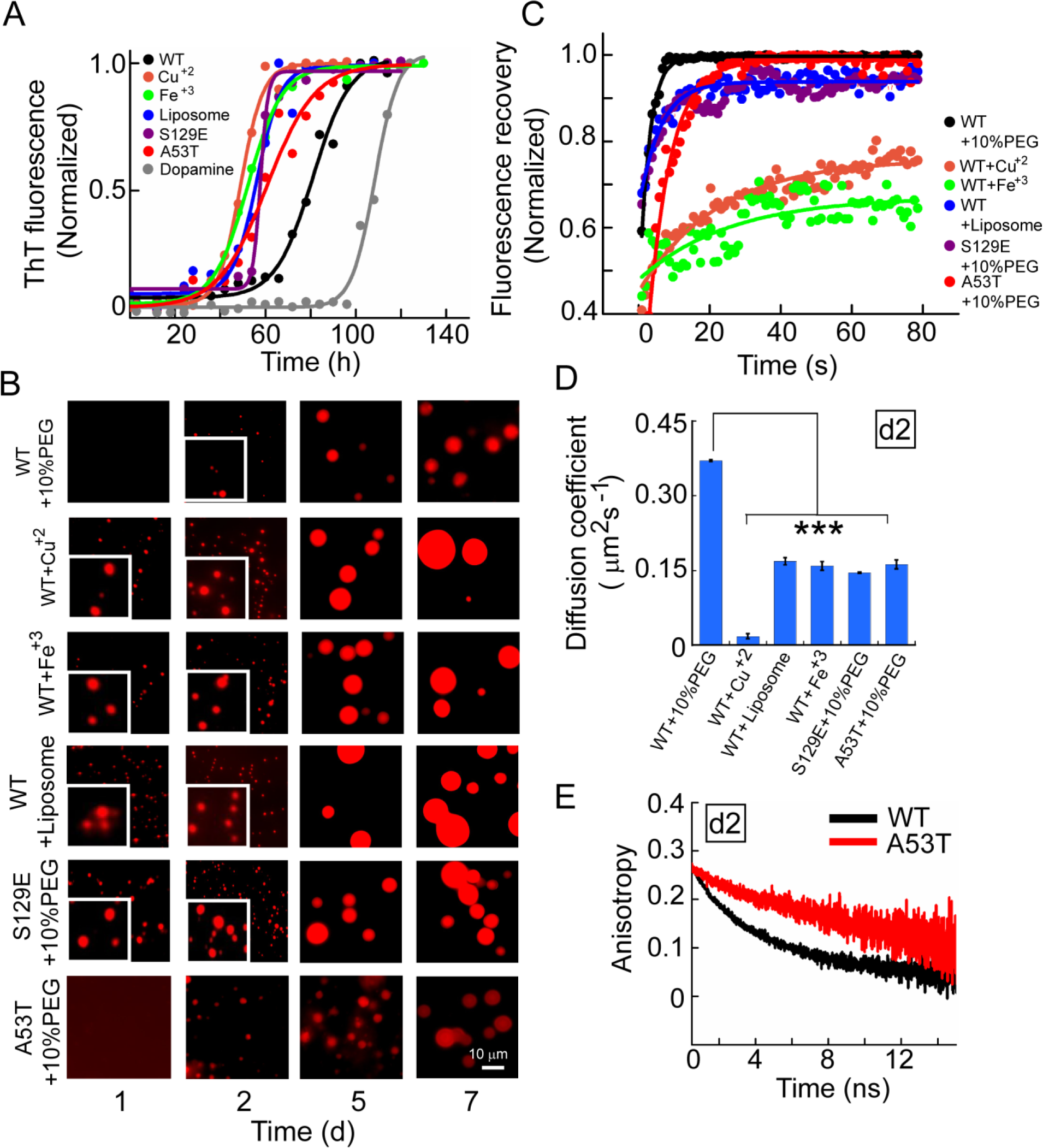
PD-associated aggregation factors promote α-Syn LLPS. **A.** ThT-fluorescence assay demonstrating α-Syn aggregation kinetics in the presence of various factors (Cu^+2^, Fe^+3^, liposomes, S129E-phosphomimetic, A53T mutation, and dopamine) is shown. 200 μM WT and mutant α-Syn were incubated with 50 μM Cu^+2^, 50 μM Fe^+3^, 1 mM liposomes and 200 μM dopamine in absence of PEG. **B**. Fluorescence microscopic images of rhodamine-labeled α-Syn droplets formed in the presence of PD-associated factors at indicated time points. The formation of liquid droplets occurred within 24 h in presence of the aggregation accelerating factors, while in the presence of PEG, the time taken was ~48 h. **C, D**. The dynamics of α-Syn molecules inside the droplets measured using FRAP. Normalized FRAP intensity curves of α-Syn droplets at day 2 (d2) (C) and corresponding diffusion coefficients (D) are shown. ω^2^ = 4 μm^2^; n=3, values represent mean ± s.e.m (***p ≤ 0.001). **E**. Time-resolved fluorescence anisotropy decay of FITC-labeled WT and A53T α-Syn from a single droplet at day 2 showing the decay is slower for A53T compared to WT protein.

We next analyzed the effect of these external factors on α-Syn dynamics inside the liquid droplets immediately after their formation. The FRAP data revealed significantly decreased fluorescence recovery for droplets formed in the presence of these additives (Figure 3C). The calculated average diffusion coefficient of the α-Syn molecules inside the liquid droplets formed in presence of Cu^+2^, Fe^+3^, and liposomes were 0.0448, 0.1592, and 0.1689 μm^2^s^−1^, respectively (Figure 3D), which is significantly less compared to PEG-induced α-Syn droplets (0.369 μm^2^s^−1^). Droplets formed by 200 μM S129E and A53T in presence of 10% PEG showed a diffusivity of 0.1457 and 0.1602 μm^2^s^−1^, respectively (Figure 3D). This is further consistent with fluorescence anisotropy decay analysis of FITC-labeled α-Syn, where A53T droplets showed slower fluorescence anisotropy decay at day 2 compared to WT suggesting more rigidity of A53T α-Syn molecules inside the droplets (Figure 3E). Our results show that although A53T mutation did not lower the critical/saturating concentration for phase separation, the physical properties such as internal viscosity or dynamics of α-Syn is greatly affected as evident by FRAP analysis. Irrespective of lower diffusivity, the average fluorescence recoveries of α-Syn droplets after photobleaching were comparable in the presence of additives, except for Fe^+3^ (62%) at day 2 (Figure S2F). The decreased diffusivity of α-Syn in the presence of these PD-associated factors could be due to enhanced aggregation leading to early maturation of the droplets. Taken together, phase separation of α-Syn correlates with its aggregation behavior in the presence of PD-associated factors suggesting that liquid droplets plausibly act as a nucleus for initiating and/or accelerating α-Syn aggregation.

### Aggregation state of α-Syn during LLPS and liquid-to-solid transition

To study the extent of aggregation of α-Syn into oligomers and fibrils during LLPS and droplet maturation, we quantified the relative amount of different species formed over time (Figure S3A). The kinetics of fibril formation measured by ThT fluorescence (inset, Figure 4A) showed that the liquid droplets are formed in the early lag phase of amyloid formation kinetics. The quantification of different α-Syn species showed mostly low molecular weight (LMW) α-Syn (~90%) along with small amount of oligomers (~8%) and fibrils (~2%) at the early stages of LLPS and aggregation (day 5) (Figure 4A). A significant decrease in LMW fraction and concomitant increase in fibril population was observed during maturation of droplets along with α-Syn aggregation. However, the amount of oligomers remained unchanged from day 5 to day 10 (Figure 4A). The increase in amyloid fibril formation over time is evident by an increase in ThT fluorescence and β-sheet structural transition monitored using circular dichroism (CD) for α-Syn solution during phase separation (Figure 4A, inset, and Figure S3B). Morphological analysis by TEM showed that fractions isolated during LLPS contain amorphous, small oligomers, and fibrillar aggregates (Figure S3C). Collectively, the data suggest that there is a dynamic interplay between the species (i.e. monomer, oligomers, and fibril); wherein, the monomer gradually converts into fibrillar aggregates via an oligomer-mediated process during LLPS.

**Figure 4:**
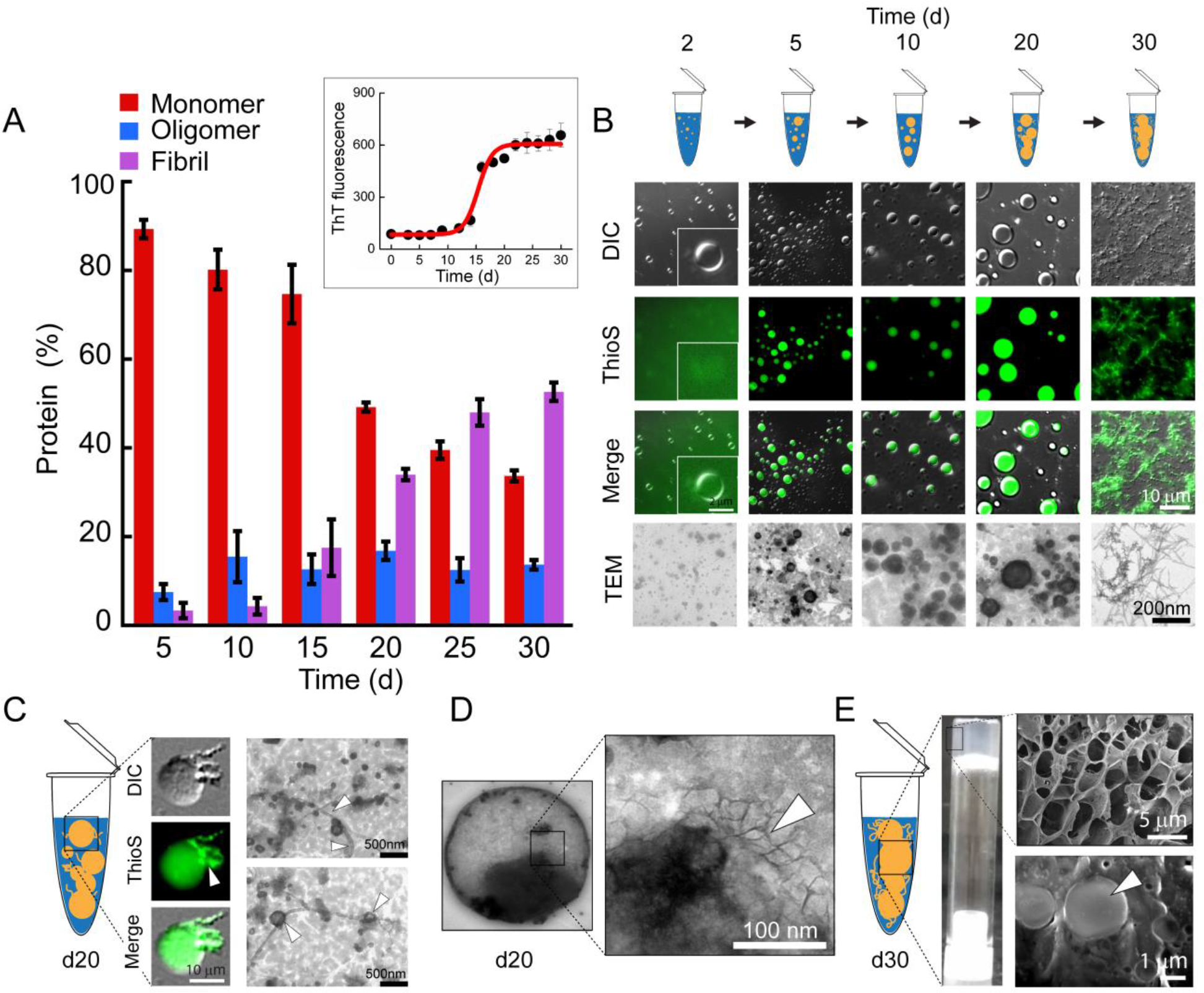
α-Syn phase separated droplets mature and age into fibrillar aggregates. **A**. α-Syn in the presence of 10% PEG in microcentrifuge tube was imaged for LLPS at various time points, and the solution was taken for isolating monomers, oligomers, and fibrils. In parallel, ThT fluorescence assay was performed to monitor the aggregation and amyloid formation by α-Syn. During LLPS and aggregation, the relative percentage of monomers (LMW), oligomers and fibrils at indicated time points were quantified and plotted. n=3; values represent mean±s.e.m. Inset shows aggregation kinetics for WT α-Syn under LLPS conditions (200 μM α-Syn + 10% PEG, no agitation, incubated at 37°C) monitored by the ThT assay. **B**. Time-dependent maturation of WT α-Syn analyzed by DIC imaging, ThioS staining, and fluorescence microscopy and TEM imaging. At day 2, α-Syn droplets showed no ThioS co-partitioning inside the droplets; probably due to lack of ThioS positive aggregates. At later time points, ThioS readily co-partitions inside the droplets, highlighting the presence of ThioS positive α-Syn aggregates during maturation of droplets. At the end of the incubation period (after 30 days), the coverslip showed a ThioS positive mesh-like structure. **C**. The Appearance of fibrils at day 20 of droplet incubation at 37°C. ThioS staining and TEM images show the presence of protein aggregates (marked by white arrows) adjacent to the droplets. **D.** Magnified TEM image showing the presence of fibril-like structures inside a droplet (white arrow). **E**. Gel formation confirmed by gel inversion test for day 30 post droplet formation by α-Syn in presence of 10% PEG at 37°C. SEM image showing the presence of liquid droplets embedded in the hydrogel bed (white arrows).

To directly establish the link between LLPS and α-Syn aggregation, we microscopically examined the liquid droplets at various time intervals after their formation in presence of 10 % PEG. ThioS staining during LLPS showed diffused fluorescence signal in the background at the beginning of the phase separation, however, it readily co-partitioned into the phase separated droplets after day 5, indicating the formation of amyloid-like species inside the droplets (Figure 4B). TEM imaging further confirmed the phase-separated α-Syn droplets formed at day 2, 5, 10 and 20, while we mostly observed fibrillar aggregates at day 30 (Figure 4B), where α-Syn liquid droplets solution transformed into a gel-like structure. Interestingly, after day 20, we observed protein aggregates emerging from the subset of liquid droplets (Figure 4C) and even fibril-like structure were observed inside the individual droplets when analyzed using high-resolution TEM (Figure 4D). As the α-Syn liquid droplet solution transformed into a gel-like state, we formally characterized its physical properties. The gel inversion test along with higher order network structure by SEM and higher storage modulus compared to loss modulus further support the gel state of the α-Syn solution (Figures 4E and S4C) (Yan and Pochan, 2010). We also asked how familial mutation, A53T, modulates the liquid-to-solid transition of α-Syn. In contrast to WT, we observed early co-partitioning of ThioS (as early as day 2) in the liquid droplets (Figure S4B), which could be due to the faster aggregation of A53T inside the droplets as compared to WT (Figure S4A). The TEM data also showed the appearance of fibrils from the A53T droplets and stiffer gel formation compared to WT α-Syn as suggested by rheological measurements (Figure S4C, inset). Overall, the data support that phase separated droplets act as nucleation sites for α-Syn aggregation and during incubation the liquid droplets harden due to the formation of amyloid fibrils.

### Domain interactions responsible for α-Syn liquid-liquid phase separation

It has been shown that multivalent IDRs and LCDs containing proteins with mostly charged amino acid residues drive the LLPS through weak intermolecular interaction (Han et al., 2012; Lin et al., 2015; Mitrea et al., 2016; Nott et al., 2015). Although, in some instances, the hydrophobic interaction mediating LLPS has also been shown (Li et al., 2018b; Pak et al., 2016). For α-Syn, the central hydrophobic NAC region primarily drives the aggregation into amyloid fibril (Giasson et al., 2001). This is strengthened by the fact that lack of the hydrophobic residue stretch in β-Syn abolishes its amyloid formation capacity (Uversky et al., 2002). To determine the domain interaction responsible for LLPS in α-Syn, we recorded 2D-HSQC-[^15^N, ^1^H]-NMR spectra for WT protein with the progression of LLPS. We also used the A53T mutant along with WT to examine potential differences. During the early stages of LLPS, WT α-Syn showed a gradual decrease in the intensity for the residues in the N-terminal along with the residues in the NAC region (Figures 5A, B). On the 20^th^ day, the peaks completely disappeared and new peaks were visible, highlighting a different aggregating α-Syn conformation (Figures S5A, C). In case of A53T, immediately after LLPS, there was a drastic decrease in the intensities (almost 60 %) for peaks at the N-termini and the NAC region highlighting the crucial role of these regions in LLPS (Figures 5A, B and S5B, D). A53T showed a comparatively greater decrease in intensities at the N-termini and the NAC region than WT after LLPS (Figure 5B). The residue-specific changes are described in detail in the Supplementary Results. The results suggest that along with IDR containing highly charged amino acids at the N-terminus, the hydrophobic NAC region is also involved in intermolecular interactions responsible for LLPS.

**Figure 5:**
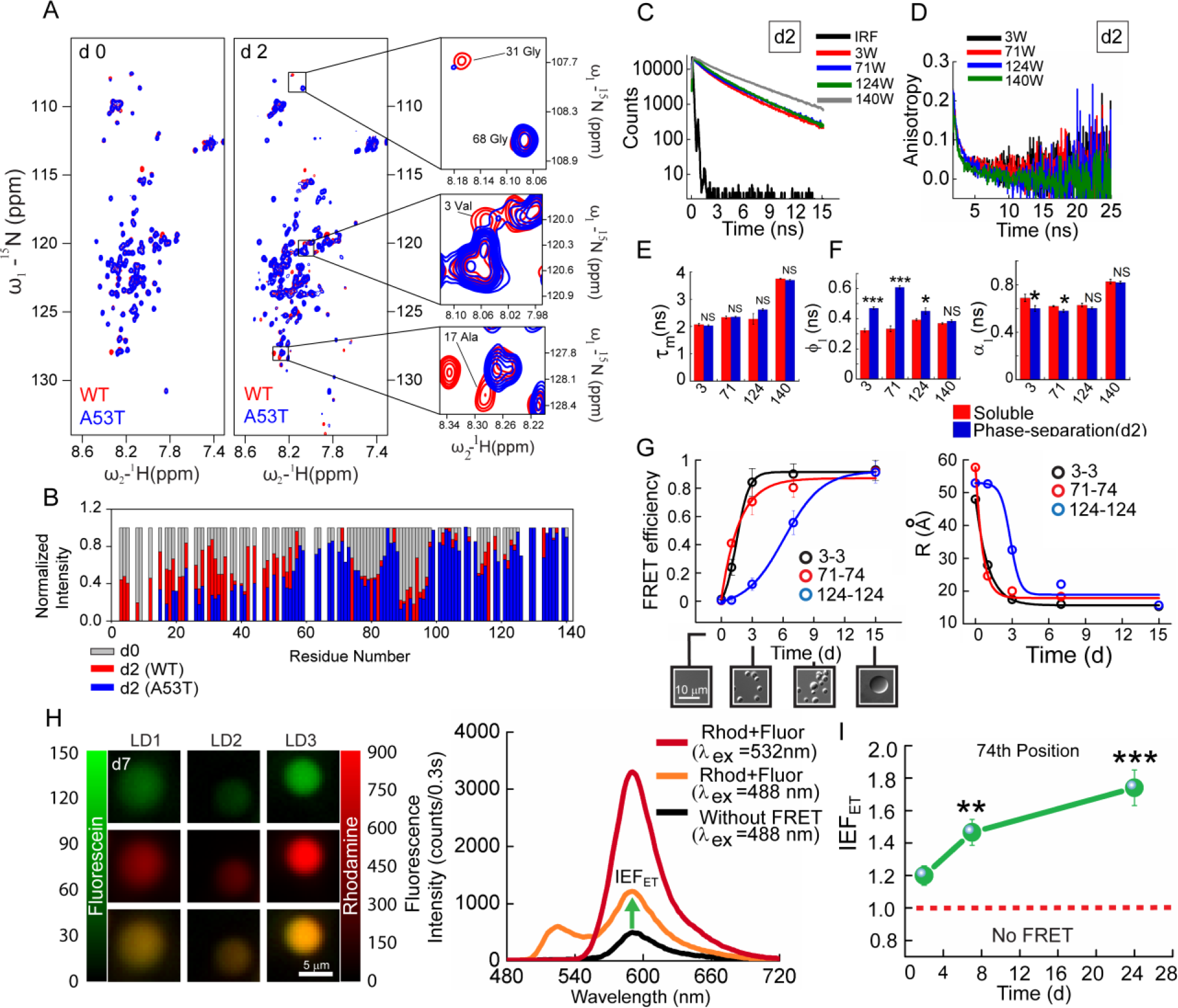
Site-specific conformational changes and dynamics of α-Syn during LLPS. **A.** Overlapped [^15^N-^1^H] HSQC spectra of WT α-Syn (red) and A53T α-Syn (blue) on day 0 and day 2 showing the residues (G31, V3, V17) of WT and A53T have significant differences in their intensities, post-LLPS. **B**. Normalized intensity (I/I_0_) profile of amide cross-peaks from ^1^H-^15^N HSQC spectra of WT (red) and A53T (blue) on day 2 (post-LLPS) showing a significant decrease in intensities for residues at N-terminus and NAC domain after LLPS. The extent of intensities decrease was more for A53T compared to WT. Time-resolved fluorescence intensity (**C**) and fluorescence anisotropy decay (**D**) of α-Syn Trp substitution mutants (positions 3^rd^, 71^st^, 124^th^, and 140^th^) post phase separation (at day 2). **E.** The mean lifetime (τ_m_) computed from time-resolved fluorescence data before (day 1, red bars) and after (day 2, blue bars) the phase separation event. **F**. The fluorescence anisotropy decay analysis reflects higher rigidity at the N terminal (3^rd^) and the NAC-region (71^st^) compared to the C-terminus. The magnitude of the first correlation time (Φ_1_) is higher for 3^rd^ and 71^st^ Trp, while the corresponding amplitudes (α_1_) are lower. **G**. *Left panel*: Spectroscopy-based FRET analysis of the bulk system involving Trp-Cys DTNB intermolecular pair. FRET efficiency for N terminal region [3W-3C (DTNB)] and the NAC domain [71W-74C (DTNB)] was higher from the beginning of the LLPS compared to C-terminus [124W-124C (DTNB)] region, which showed higher FRET during the later phase of liquid-solid transition. *Right panel*: The intermolecular distance (R) calculated from the FRET efficiency values were plotted against time considering the R_0_ value as of 23 Å. **H**. *Left*: Single droplet fluorescence imaging of the droplets containing fluorescein (donor) and rhodamine (acceptor) as intermolecular FRET pairs for the 74^th^ position (NAC). *Right*: Red and orange lines show emission spectrum from a droplet excited at the 532 nm and 488 nm, respectively. To note, rhodamine emission at 488 nm excitation shows enhanced emission compared to without energy transfer scenario (black line) from the same droplet indicating the closeness of the fluorophore due to energy transfer. **I**. The evolution of the energy transfer at the 74^th^ position due to intermolecular FRET showed that the NAC regions of α-Syn come close to each other with time during aggrgeation inside the droplets. n=3, values represent mean ± s.e.m (**p ≤ 0.005; ***p ≤ 0.001).

To further delineate the relative involvement of domain interaction during LLPS and liquid-to-solid transition, we strategically introduced single Trp probes (α-Syn is devoid of Trp) at the N-terminal (3W), NAC (71W) and C-terminal (124W, 140W) domains and subjected the protein to undergo LLPS. Previous studies showed that this introduction of Trp did not majorly alter the properties of the protein (Sahay et al., 2015). To investigate the difference in the local environment and structural rigidity at these four different positions, we employed time-resolved fluorescence intensity and fluorescence anisotropy decay analysis for day 2-old LLPS sample (immediately after droplet formation) and a sample without LLPS. The fluorescence intensity and anisotropy decay curve for the different Trp probes pre and post LLPS are given in Figures 5C, D and S6A. The data show that although there is no significant change in the local environment of the Trp probes representing each domain of α-Syn (Figure 5E), the structural rigidity of the N-terminus and the NAC region (3^rd^ and 71^st^ positions, respectively) is considerably high compared to the C-terminus (Figure 5F). The rigidity is presented by the faster correlation time (Φ_1_) and its amplitude (α_1_) (Sahay et al., 2015). The Trp probe at 3^rd^ and 71^st^ positions showed higher values of Φ_1_ and lesser values of α_1_, which clearly indicates increased rigidity at these positions post-LLPS (Figure 5F).

Subsequently, we asked whether the structural rigidity at the N-terminus and the NAC region is due to the self-association of the protein molecules after phase separation. To address this, we introduced single cysteine residue at positions; N-terminal (3C), NAC (74C) and C-terminal (124C). Next, we labeled the cysteine mutants with DTNB (a non-fluorescent acceptor), and performed Trp-DTNB intermolecular FRET with complimentary Trp (donor) and Cys-DTNB (acceptor) pairs (Jha and Udgaonkar, 2009). FRET was quantified by the decrease in fluorescence intensity by Trp. The FRET efficiency for the N-terminal (3W-3C) and the NAC (71W-74C) regions increased rapidly after phase separation at day 2. The energy transfer for C-terminus (124W-124C), however, was low compared to the N-terminal and NAC regions (Figures 5G and S6B). The distance between the FRET pairs was also calculated for each of the positions considering the R_0_ value as 23 Å (Bhatia et al., 2018), which reflects increased proximity and association among these domains over the time (Figure 5G). The fluorescence spectroscopy data reveals that during the early stages of LLPS, the inter-domain interaction between N-terminus and NAC region might drive the formation of the condensed protein-rich phase separated droplets. Important to note that all the above experiments probing domain interaction were ensemble measurements reflecting properties of α-Syn inside and outside the droplet.

To confirm the domain involvement at a single droplet resolution, we studied the FRET using α-Syn NAC domain as a representative case. For this, fluorescein-5-maleimide-labeled α-Syn at 74^th^ position was chosen as the donor and rhodamine-C2-maleimide labeled α-Syn at the same position was chosen to be the acceptor fluorophore. We prepared three types of phase-separated samples; droplets containing only donor (fluorescein-α-Syn); only acceptor (rhodamine-α-Syn); and droplets containing an equimolar ratio of both donor and acceptor. The three samples were incubated at 37°C so that they undergo LLPS. When the donor fluorophore was excited, in the sample containing donor and acceptor α-Syn in an equimolar ratio, the fluorescence emission signal from acceptor fluorophore was detected indicating intermolecular FRET (Figure 5H). No energy transfer was observed in only fluorescein and rhodamine controls (data not shown). We further estimated the efficiency of FRET with time, and our data showed that during incubation, the extent of FRET gradually increased (Figures 5I and S6C). This further supports the ensemble measurement that NAC region is actively involved in LLPS and droplet maturation. The detailed FRET results and experimental set-up are given in Supplementary Information.

To gain further insight into the role of the individual domains in LLPS, we probed human β-synuclein (β-Syn, lacking 8 residues hydrophobic amino acid stretch in the central NAC domain amongst others), γ-synuclein (γ-Syn) and the amyloid core α-Syn (30-110) comprising residues 30-110 for liquid droplet formation. β-Syn did not show amyloid formation at physiological pH, while γ-Syn shows extremely slow aggregation *in vitro* (Uversky et al., 2002). In contrast, the α-Syn core (30-110) shows faster aggregation into fibrils compared to the full-length protein (Vilar et al., 2008). As anticipated, ThT fluorescence assay showed faster aggregation kinetics for α-Syn core (30-110), whereas β-and γ-Syn did not show any considerable aggregation even after 120 h of incubation at pH 7.4 (Figure S4D). When we examined the LLPS behavior of these proteins (200 μM of protein and 10% PEG), α-Syn core (30-110) (lacking N-terminus and C-terminus) underwent LLPS at a faster rate (within 24 h) compared to full-length α-Syn (48 h), while β-or γ-Syn did not phase separate even after 6 days (Figure S4E). Prolonged incubation in LLPS conditions led to amyloid fibril formation for α-Syn core (30-110), similar to WT as confirmed by TEM (Figure S4F). This suggests that the hydrophobic patch in the core region of α-Syn is primarily essential both for LLPS and amyloid aggregation.

### Liquid-liquid phase separation and liquid-to-solid transition of α-Syn in mammalian cells

Since α-Syn undergoes LLPS *in vitro*, we hypothesized that it might also phase separate in cells under appropriate conditions. In order to assess the behavior of α-Syn in a cellular environment, we used HeLa cells stably expressing FLAG and tetracysteine-tagged α-Syn (C4-α-Syn) because of its robustness, controlled expression upon doxycycline induction, and widely used model cell line for studying *in cell* LLPS (Maharana et al., 2018; Maucuer et al., 2018; Molliex et al., 2015). The small (∼1.5 kDa) tetracysteine-tag which binds to biarsenical compounds such as FlAsH has been used to monitor protein dynamics in live cells (Irtegun et al., 2011; Whitt and Mire, 2011). *In vitro* and *in cell* characterization of C4-α-Syn suggests that it shares identical biochemical and biophysical properties with WT α-Syn (Roberti et al., 2007), thus providing a suitable system for monitoring α-Syn LLPS in cells.

To perform live cell imaging experiments, we cultured the stably transfected HeLa cells for 24 and 48 h and subsequently stained C4-α-Syn with FlAsH. We found that ~20 % of the cells showed cytoplasmic protein accumulates (Figure 6A), whereas remaining cells showed diffused pan-cellular localization of C4-α-Syn. The background fluorescence for FlAsH was estimated with two controls, one with non-α-Syn expressing HeLa cells and other with pcDNA-α-Syn transfected cells, which showed notably lower signal compared to cells expressing C4-α-Syn (data not shown). Several reports suggest that treatment of cells with metal ions, such as Fe^+3^ induce reactive oxygen species (ROS) formation and promote aggregation of α-Syn in cells that over-express the protein (Matsuzaki et al., 2004; Ostrerova-Golts et al., 2000). Therefore, we examined the effect of iron on the aggregation and phase separation behavior of α-Syn in cells. Strikingly, post 24 h of treatment with 10 mM of ferric ammonium citrate, >95 % of cells displayed cytoplasmic droplet-like assemblies (235 ± 20 droplets/cell) (Figures 6A and S7A, B). Confocal microscopy revealed that these assemblies were majorly spherical with an average diameter of 0.46 μm (Figure 6B). However, to note, after 48 h of iron treatment, these droplets were largely clustered and localized majorly at the perinuclear region of the cell (Figures 6A and S7A). Furthermore, the number of the droplets decreased considerably after 48 h (76 ± 21 droplets/cell) (Figure S7B) with the increase in the average diameter of the droplet (Figure 6B, left panel).

**Figure 6:**
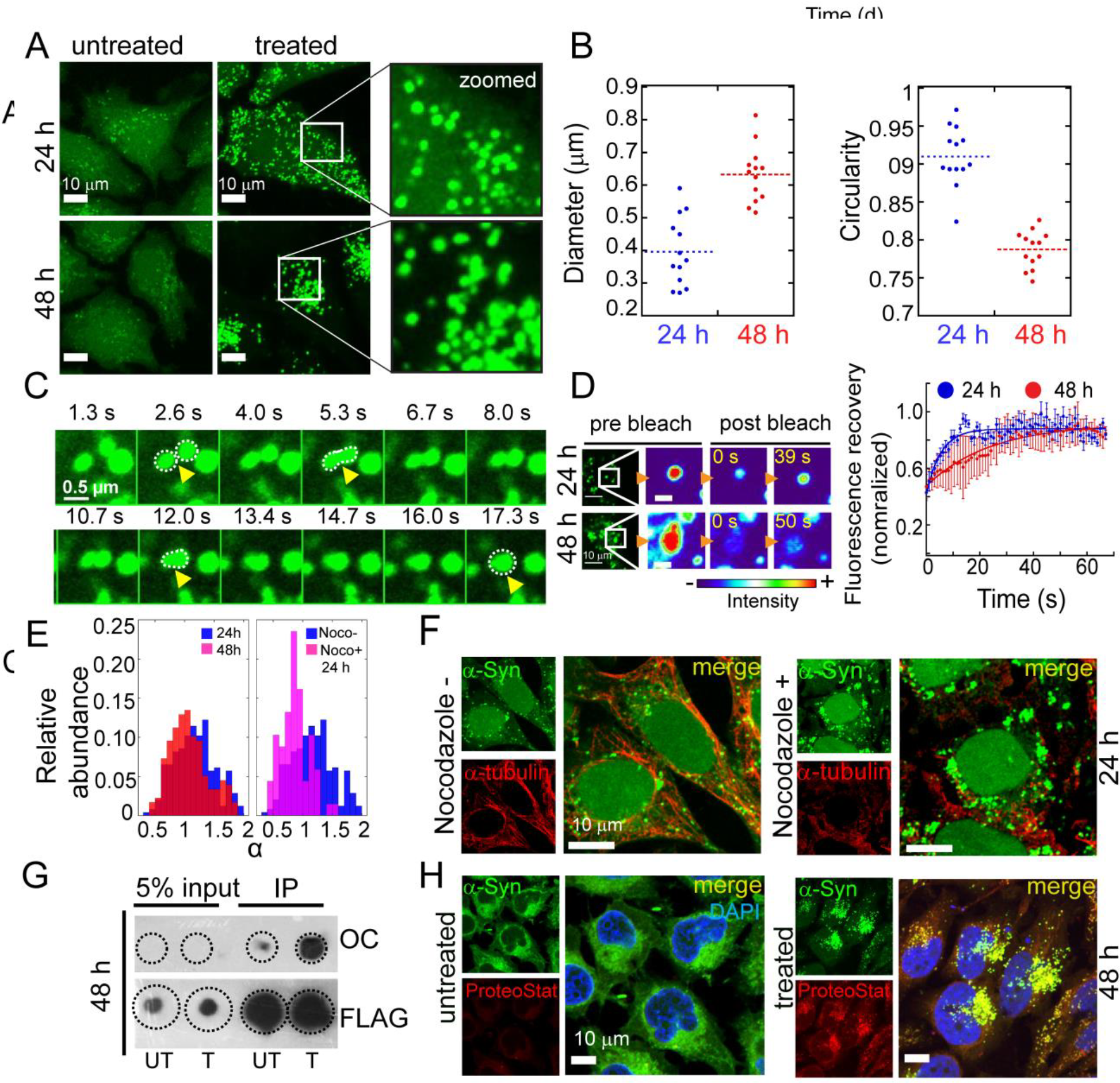
Liquid-like condensates of α-Syn *in cells*. **A**. Confocal images of HeLa cells over-expressing C4-α-Syn stained with FlAsH-EDT2 captured at 24 h and 48 h. Untreated cells predominantly show pan-cellular localization; however, cells treated with 10 mM ammonium ferric citrate showing intracellular liquid-like droplet formation. Scale bar: 10 μm. **B**. Quantification of the diameter and circularity of droplets in iron-treated cells. The number of images processed form from three independent experiments are n>12, and the total number of droplets accounted is n = 2400 (for 24 h) and n = 950 (for 48 h). Data is presented as a dot plot with the corresponding mean represented as dash lines. **C**. Images represent a time-lapse series of the fusion event. Scale bar: 0.5 μm. **D**. *In cell* FRAP recovery of droplets at 24 h and 48 h of treatment. Left panel: The images correspond to the region of interest (ROI) with pre-bleach and post-bleach droplets at t=0 s and t=39 s (24 h) and t = 50 s (48 h). Right panel: Normalized fluorescence recovery curves for 24 h and 48 h droplets showing slow recovery for 48 h droplets compared to 24 h. Data are shown as mean ± s.e.m (n=5). Due to the difference in the bleach ROI, diffusion constant could not be computed. Scale bar: 1 μm. **E.** Distribution plot of the diffusion exponent (α) calculated based on the log-log fit of mean squared displacement versus time. The plots present a comparison of droplet behavior after 24 h and 48 h of iron treatment (left) and 24 h with and without treatment of nocodazole (right). **F**. Immunofluorescence images for α-tubulin staining of 24 h iron-treated HeLa cells with or without nocodazole. α-Syn droplets were clumped upon nocodazole treatment. Scale bar represents 10 μm. **G.** Immunoprecipitation of α-Syn using anti-FLAG from cell extracts of iron-treated and untreated cells, post 48 h, and subsequent dot blots probed with amyloid-specific OC antibodies. Significantly increased OC staining was observed for treated samples compared to untreated cells showing higher fibril formation due to LLPS. Samples were probed with FLAG antibody to ensure equal loading of the immunoprecipitates (IP). UT and T indicate untreated and treated with iron. 5% input of the total protein is also shown. **H.** Aggresome detection using ProteoStat-dye staining. Post 48 h treatment, α-Syn localized at the perinuclear region showing colocalization with the ProteoStat dye, while the untreated cells show no dye binding.

To probe the dynamic behavior α-Syn inside the droplets, we performed time-lapse confocal microscopy of cells treated with iron at different time points. At 24 h, we observed that these droplets are highly mobile, and undergo frequent events of fusion with rapidly relaxing back into spherical assemblies (Figure 6C). These α-Syn droplets formed in the cell showed characteristic properties of liquid-state, which includes spherical morphology and fusion ability (Figures 6B and C), thereby suggesting that these accumulates are liquid-like droplets formed by α-Syn LLPS in the cytoplasm. Furthermore, they did not show colocalization with intracellular lipid droplets (detected using Nile red staining (Greenspan et al., 1985) and membrane-bound organelles such as lysosomes, mitochondria (Figure S7C), thus confirming its liquid-like membrane-less state.

In order to understand the dynamic properties of C4-α-Syn molecules in these droplets, we performed *in cell* FRAP measurements, post 24 and 48 h of stressor treatment. As a control, the region with diffused expression was chosen to determine the intracellular dynamics of the FlAsH stained C4-α-Syn molecules. The recovery for the droplet-free region was partial; with percent recovery of ~56 % and t_1/2_ ~ 10s, suggesting that the fluorescently labeled C4-α-Syn slowly diffuses in the cellular milieu (data not shown). Post 24 h treatment, the fluorescence recovery in the bleached region of the droplet was estimated to be 57 ± 16 %, and recovery half-time of ~5.4 s (Figure 6D), indicating that α-Syn molecules can diffuse and equilibrate with the cytoplasmic pool, in agreement with the properties of the liquid-like state. In contrast, at 48 h, the % recovery reduced to ~36 %, with a significant increase in recovery half time (t_1/2_~23 s) (Figure 6D). The decreased mobility of α-Syn molecules in these droplets can probably be attributed to the liquid to solid-like transition, which might be due to the aggregation of C4-α-Syn within the droplets.

Further, single particle tracking and statistical analysis were done to understand and track droplet motion inside the cells. More than 400 droplets in cells across different conditions were taken into account for the analyses. The time-dependent trajectory data was generated using IMARIS software and analyzed with *msdanalyser* package in MATLAB2018B (Tarantino et al., 2014). At 24 h and 48 h, the mean squared displacement of the droplets as a function of time was computed (Figure S7D) and the diffusive exponent (α) was determined. The major population of the droplets at 24 h showed α values to be greater than 1, among which few have α values close to 2 (Figure 6E), indicating a super-diffusive, nearly directed motion (Munder et al., 2016; Tarantino et al., 2014). This suggests that the motion of the droplets, at 24 h, possibly involves an active or facilitated movement. The representative trajectories of single droplets at 24 h for different values of α are provided in Figure S7E. Remarkably, after 48 h, a population shift in the α-distribution was observed where the mean was around α≤1 (Figure 6E, left panel), indicating diffusive or sub-diffusive behavior of these droplets. It is important to note that droplets formed after 24 h are randomly distributed in the cytosol, while they cluster at the perinuclear region post 48 h treatment (Video S2 and S3). This indicates that possibly the particles are initially actively displaced and directed towards the perinuclear site, and later its motion is influenced by the size and particles’ microenvironment. To probe the mechanism of super-diffusive displacement of α-Syn droplets and the possible involvement of microtubule for this, we treated the cells with nocodazole to de-polymerize the microtubules that might be assisting the active movement of the droplets. Interestingly, upon microtubule destabilization confirmed using α-tubulin staining, the majority of the droplets formed at 24 h clumped together (Figure 6F, right panel, and Video S4) suggesting that microtubule network might play a role in droplet movement. When we calculated α-distribution post microtubule destabilization, we observed a considerable shift towards values equal to or less than 1, indicating a freely diffusive motion (Figure 6E, right panel) further supporting the role of the microtubule network for droplets movement. We further determined the track straightness value, which is the ratio of displacement and length of the path traveled by the particles. The increased value of this ratio denotes a more directed and confined motion of the particle (Fusco et al., 2003). For both 48 h and nocodazole-treated 24 h cells, the distribution of straightness weighted towards lower values compared to 24 h cells (Figure S7F). The track straightness of the droplets at 24 h also showed a positive correlation with α (Figure S7G), further supporting the directed motion of these droplets. In addition, distribution of the root mean square velocity (V_rms_) for the droplets showed that at 48 h or in the nocodazole-treated cells, the droplets V_rms_ was lesser compared to droplets formed at 24 h (Figure S7H). This analysis corroborates with an increase in the effective size of the droplets at later time-point or clumping upon nocodazole treatment. The results suggest that α-Syn liquid droplet movements are much more directed with the assistance of the microtubules, which is significantly reduced after their liquid-to-solid transition and its localization to the perinuclear area.

### Liquid-to-solid transition of α-Syn triggers aggresome formation

Since α-Syn liquid droplets in cell harden with time as suggested by FRAP data, we asked whether these droplets trigger the α-Syn aggregation with production of fibrils in cells. Previous studies have reported that PD-associated factors, including iron, promote α-Syn aggregation in the transfected cells such as COS-7 and BE-M17 cells (Lee et al., 2002; Paxinou et al., 2001). To test this, we probed α-Syn amyloid-like fibril formation using the fibril-specific OC antibody (Kayed et al., 2007) staining with immunoprecipitated α-Syn at different time-points. Dot blot assay revealed that the OC signal increased for both treated and untreated cells over time (Figures 6G and S8A). However, 48 h treated cells showed considerably increased immunoreactivity for OC compared to the untreated cells (Figure 6G). The samples were probed with anti-FLAG to ensure equal loading of the immunoprecipitates. The dot blot result indicates the gradual transition of liquid droplets into solid-like state consisting of amyloid-like aggregates. We further tested cellular viability to examine the effect of liquid-to-solid transition on cellular processes. We found that the cell viability remained uncompromised even in the presence of α-Syn aggregates (Figure S8B), which is in contrast to neuronal cells where iron-induced α-Syn accumulation and aggregation leads to neurotoxicity (Yasuda et al., 2013). The perinuclear localization of α-Syn droplets after 48 h and lack of cellular toxicity suggest that α-Syn droplets might form aggresome, which is known to provide cytoprotectivity against unwanted protein aggregates (Garcia-Mata et al., 1997). To probe this, we performed immunofluorescence and quantitative flow cytometric analysis with the aggresome specific marker ProteoStat. The colocalization of α-Syn and ProteoStat was monitored, and aggresome propensity factor was calculated for different time-points (Figures 6H and S8C, D) after iron treatment. The data clearly revealed that iron-induced LLPS triggers α-Syn aggresome formation in HeLa cells after 48 h and possibly facilitate their clearance by a yet unknown mechanism.

## DISCUSSION

Although the pathway(s) and mechanism of α-Syn aggregation in PD have been studied extensively, the early events that lead to toxic α-Syn aggregates remain unclear. In this study, we elucidate the detailed mechanism of LLPS driven α-Syn aggregation and address fundamental questions regarding the formation of amyloid aggregates in Parkinson’s disease.

α-Syn is an intrinsically disordered protein with two LCDs, one present at the N-terminal region (residues 10-23) and another at the NAC region (residues 63-78) (Figure 1A). We hypothesized that α-Syn aggregation involves LLPS and liquid to solid-like transition as shown by other IDR-containing proteins like FUS and TDP43 (Conicella et al., 2016; Molliex et al., 2015; Patel et al., 2015). Indeed, our data show that α-Syn undergoes LLPS in the presence of PEG (Figure 1B), suggesting PEG-induced molecular crowding increases the local concentration and protein association (Shtilerman et al., 2002) thereby resulting in LLPS. Increase in the α-Syn intermolecular interactions by lowering pH (pH 5.4) (Wu et al., 2009) also promotes α-Syn phase separation with a significant decrease in the solubility limit (5 μM concentration, Figure 1E). Initially, α-Syn droplets possess liquid-like properties such as fusion and spherical shape (Video S1), rapid internal molecular rearrangements (high FRAP recovery) (Figure 2C), and temperature reversibility (Figure 1F), similar to other phase separating proteins (Lin et al., 2015; Molliex et al., 2015; Patel et al., 2015). However, over time, the droplets become more rigid with less dynamic α-Syn molecules, consistent with a liquid to solid-like transition, as observed for FUS, TDP 43 (associated with amyotrophic lateral sclerosis) (Conicella et al., 2016; Molliex et al., 2015; Patel et al., 2015), and Tau (associated with Alzheimer’s disease) (Ambadipudi et al., 2017; Wegmann et al., 2018).

LLPS of α-Syn could be strongly implicated in PD, as various PD-associated conditions such as exposure to metal ions (i.e. Fe^3+^ and Cu^2+^), lipids, the phosphomimetic substitution at serine 129, and a familial mutation (Breydo et al., 2012) promote LLPS by lowering the solubility limit of α-Syn and increase in the internal rigidity of liquid droplets (Figures 3B and C). In contrast, conditions that delay synuclein aggregation (such as the β- and γ-Syn), (Figures 3A and S4D), slow down the LLPS significantly (Figures S2C and 4E). These correlative findings suggest that phase separation and aggregation are interlinked, where LLPS is a critical step for aggregation of α-Syn. This is further strengthened by the finding that liquid droplets first appear in the early lag phase of α-Syn aggregation and the presence of fibrils inside the droplets in the elongation phase (Figures 4B and D). α-Syn remains mostly in the monomeric state during early stages of LLPS (Figure 4A), as weak intermolecular interactions by protein govern the liquid droplet formation (Banani et al., 2017; Shin and Brangwynne, 2017). However, over time, monomeric α-Syn gradually changes to oligomers and fibrils, suggesting that oligomers and fibrils occur concurrently with the liquid to solid-like transition. This is further evident from the increase in size, change in droplet morphology, the formation of ThioS positive aggregates and visible fibrillar aggregates within these droplets during the transition (Figure 4). Our observations are consistent with several previous studies including Tau (Ambadipudi et al., 2017; Wegmann et al., 2018), TDP 43 (Conicella et al., 2016), FUS (Molliex et al., 2015; Murakami et al., 2015; Patel et al., 2015), which are shown to undergo an irreversible transition of liquid-droplets to protein aggregates.

After long incubation (for a month), the phase separated α-Syn solution eventually convert to a gel state (Figures 4E and S4C). Our previous data suggest that the formation of an amyloid hydrogel primarily serves as a reservoir for cytotoxic α-Syn oligomers and fibrils (Kumar et al., 2018). This data indicates that LLPS and gel formation by α-Syn could be a toxic process associated with PD. Other proteins such as Tau, FUS, TDP43, and hnRNPA1 protein have also been shown to form protein-rich droplets and eventually convert to a gel state during aggregation (Kato et al., 2012; Molliex et al., 2015; Monahan et al., 2017; Murakami et al., 2015; Wegmann et al., 2018). However, it is still unclear whether LLPS is essential for α-Syn aggregation into oligomers and fibrils or other pathways (exclusive of LLPS) could lead to α-Syn fibril formation.

It is known that weak non-specific interactions through IDRs and LCDs of protein mediate the LLPS, however more specific interactions like interaction between β-strands forming fibrils might facilitate the liquid to solid/gel state transition (Alberti, 2017; Banani et al., 2017; Boeynaems et al., 2018; Hyman et al., 2014; Shin and Brangwynne, 2017). Although α-Syn is known to aggregate primarily through hydrophobic interactions mediated by the NAC region (Giasson et al., 2001), our previous studies showed that early oligomerization events require interaction through the N-terminus (Sahay et al., 2015). This parallels with the presented NMR and specific FRET analysis of LLPS, which shows that both N-terminus as well as the middle NAC regions are involved in LLPS (Figure 5). Further, the experiments show that for liquid-to-solid transition, the C-terminus is also involved (Figure 5G). However, for α-Syn liquid droplet formation, N-terminus containing IDRs and LCDs are not essential as the core domain of α-Syn (i.e. α-Syn (30-110)) exhibited rapid LLPS. This suggests that the hydrophobic interaction could alone drive LLPS and liquid-to-solid transition, similar to fibril formation (Winner et al., 2011). The absence of hydrophobic interaction mediated by NAC region could abolish the LLPS of α-Syn, as synuclein lacking the middle hydrophobic stretch, i.e. β-Syn, did not form any droplets even after prolong incubation (Figure S4E).

Previous studies showed that at high local concentration, or change in pH, temperature or access with specific ligand binding (such as RNA), proteins undergo LLPS in cells, which could be reversible and tightly controlled (Maharana et al., 2018; Munder et al., 2016; Nott et al., 2015; Shin et al., 2017). In contrast, proteins associated with neurodegenerative orders, liquid-to-solid transition along with fibril formation was also seen (Kwon et al., 2013; Molliex et al., 2015; Wegmann et al., 2018). α-Syn under iron-induced oxidative stress (Matsuzaki et al., 2004; Ostrerova-Golts et al., 2000) readily forms liquid-like droplets with rapid turnover and fusion ability (Figure 6C, D). With time these liquid droplets, however, change to solid-like state with the formation of OC antibody-positive amyloid-like aggregates (Figure 6G). Further, the liquid-to-solid transition of α-Syn droplets leads to aggresome formation with no cytotoxicity (Figures 6H and S8B). Apart from the larger size, the perinuclear α-Syn condensates also show restricted movement compared to liquid droplets formed at an earlier time point (24 h). Since aggresome formation actively requires microtubules (Garcia-Mata et al., 1999; Johnston et al., 2002), we hypothesized that microtubules might help in the α-Syn liquid droplet movement and aggresome formation. Indeed, microtubule destabilization by nocodazole in cells halted droplet movement and clustered the droplets into large aggregates (Figure 6E and F). The data indicate that microtubules possibly assist in the droplet movement and their maturation into aggresomes. In this context, it has been shown that actin filaments influence the localization and movement of liquid condensate of LAT cluster (Kaizuka et al., 2007; Yi et al., 2012). The F-actin scaffold is also shown to provide mechanical support to the large nucleus of *Xenopus laevis* oocytes against gravitational forces (Feric and Brangwynne, 2013).

The concept of intracellular phase separation and formation of the toxic membrane-less compartment is an emerging concept in the field of protein aggregation in neurodegenerative diseases (Aguzzi and Altmeyer, 2016; Brangwynne et al., 2009; Brangwynne et al., 2011). Based on our observations, we propose a model for α-Syn aggregation, which is initiated through liquid-liquid phase separation (Figure 7). Under physiological conditions, α-Syn does not undergo phase separation, possibly due to its auto-inhibited conformational state (Dedmon et al., 2005), but at high local concentrations; due to metal exposure, lipid interaction, pH changes, the protein phase separates and forms liquid droplets. Similarly, in cells, upon cellular insult such as oxidative stress; soluble disordered α-Syn phase separates to form dynamic liquid-like droplets. The protein-dense droplets are stabilized by weak intermolecular interactions involving residues at the N-terminus and NAC domain leading to the high local concentration of α-Syn, thus, providing a nucleus for aggregation. Slow maturation and aging of the droplets eventually convert it into hydrogels composed of structurally ordered oligomers and amyloid fibrils.

**Figure 7:**
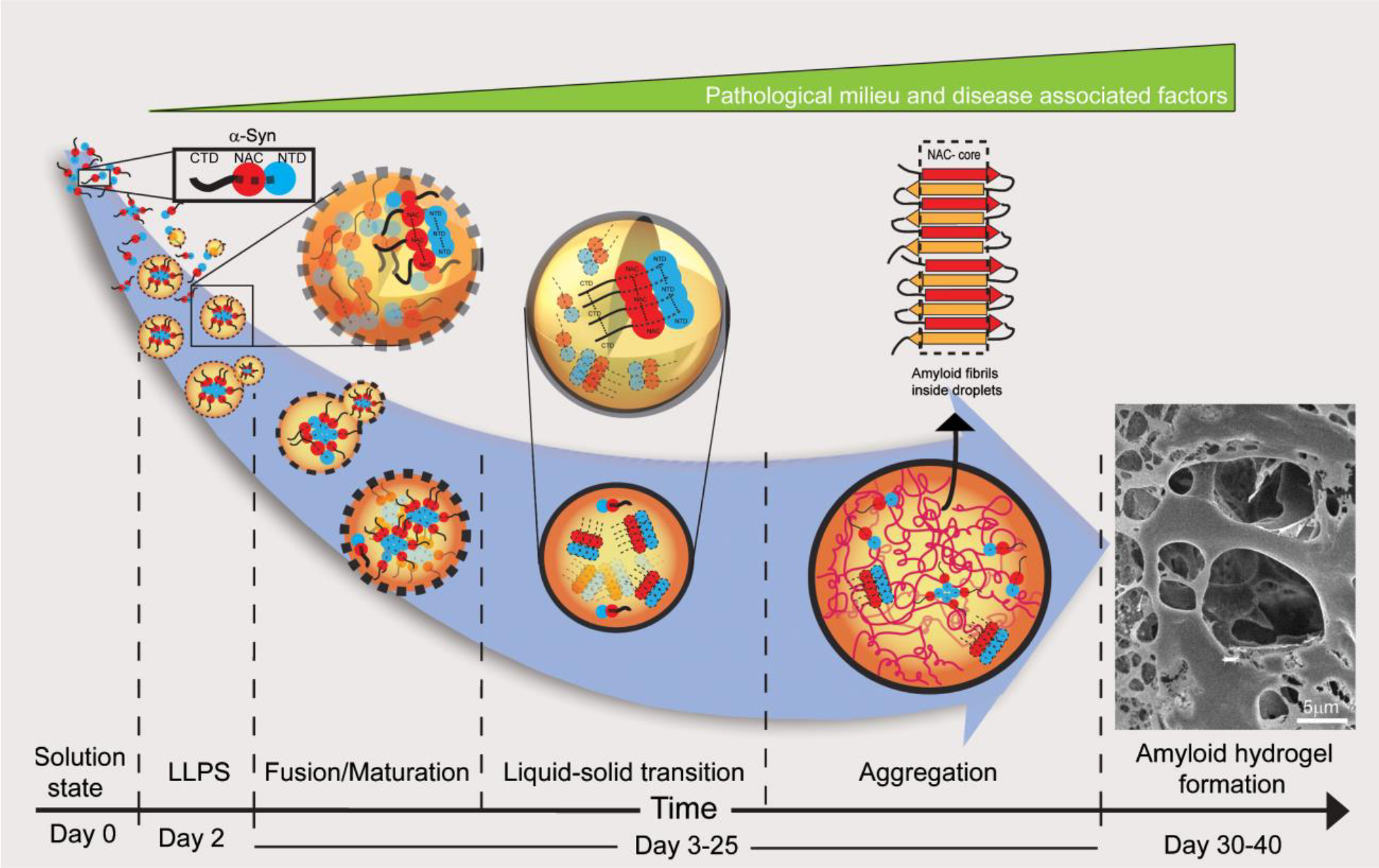
Schematic representation of the proposed mechanism for α-Syn LLPS and aggregation. Monomeric α-Syn can undergo a phase transition event in the pathologically relevant milieu and in the presence of disease-associated factors. The phase separated droplets mature from a liquid state to a solid-like state due to fusion and Ostwald ripening. α-Syn molecules inside these droplets gradually become stiffer and eventually transform into amyloid hydrogel state containing fibrillar aggregates and oligomers. The intermolecular interactions involving the residues in the N-terminal and NAC domains primarily drive the liquid-to-solid-like transition of α-Syn.

In summary, the present study provides new insights into the mechanism of initiation and progression of α-Syn aggregation through LLPS. More importantly, it provides a detailed map of the pathophysiological conditions that can promote LLPS and subsequently α-Syn aggregation, thus contributing towards understanding the processes involved in PD pathogenesis. Delineating the process of phase separation and progressive aggregation could help, particularly, in understanding its function and malfunction in protein aggregation-associated neurodegenerative diseases.

## EXPERIMENTAL PROCEDURES

### Expression and purification of recombinant synuclein proteins

α-Syn WT and mutants including A53T, tryptophan (3W, 71W, 124W) and cysteine variants (3C, 74C, 124C), phosphomimetic S129E α-Syn, core α-Syn (residues 30-110), β-and γ-Syn proteins were expressed and purified using previously reported protocols (Volles et al., 2007) with slight modifications (Singh et al., 2013). For the Cys substituted mutants (3C, 74C, 124C) of α-Syn, the protein was purified in a reducing environment (1 mM DTT) to prevent covalent disulfide bonding between the intermolecular Cys residue, which might have led to dimerization. Proteins were labeled with fluorescein, rhodamine dyes, and DTNB as per the manufacturer’s instructions (ThermoFisher Scientific, USA). The detailed expression and purification process are described in the supplementary information.

### *In-vitro* liquid-liquid phase separation

To study LLPS and liquid droplet formation, the proteins of interest (α-Syn and its variants) were suspended in 20 mM phosphate buffer (pH 7.4), 0.01% sodium azide. 10 μl of the reaction mixture prepared under various LLPS screening conditions; including PEG-8000, Cu^+2^, Fe^+3^, liposomes, dopamine at mentioned concentrations were drop-casted onto glass slides and sandwiched with a 12 mm coverslip (Blue Star, India). The slides were kept in a moist chamber (37°C) for subsequent incubation periods and further visualized with 63X oil immersion objective under a DMi8 microscope (Leica, Wetzlar, Germany) in the DIC mode.

### Fluorescence recovery after photobleaching (FRAP) experiment

WT and A53T α-Syn were labeled using NHS-rhodamine C2 (ThermoFisher Scientific, USA) as per the manufacturer’s instructions. For *in vitro* FRAP study, LLPS samples were spotted onto 15 mm depression slides (Blue Star, India). Photo-bleaching of the phase separated droplets was carried out using a 561 nm DPSS 561-10 laser at the 5^th^ cycle and recovery was recorded till the fluorescence emission reached a plateau. Appropriate ROIs outside and inside the droplet (adjacent to bleached region) were taken as controls. FRAP studies were performed using a laser scanning confocal microscope (Zeiss Axio-Observer Z1 microscope) with a 63X oil immersion objective. The selected region of interest (ROI) was bleached with a 100 % laser power. All the measurements were performed at room temperature and at least in triplicates. For *in cell* FRAP, the bleaching of ROI (2-3 μm) was performed at a 100% laser power and 50 iterations, after acquiring 2 images before bleaching and the recovery was monitored for ~100 s. The images were corrected for laser bleaching by selecting a fluorescent region outside the ROI. FRAP data analysis was performed as described (Supplementary information).

### Time-Correlated Single Photon Counting (TCSPC) microscopy

To determine the fluorescence anisotropy decay and lifetime decay analysis, WT and A53T α-Syn were labeled with FITC as per the manufacturer’s protocol (ThermoFisher Scientific, USA). Fluorescence anisotropy decay curve of FITC-labeled proteins inside a single droplet was acquired with a time-resolved microscopy setup (MT-200, PicoQuant®, Berlin). The 482 nm laser (at 5-8 μW power) pulsing at 40 MHz, and a dual-band dichroic 480/645 (Chroma) was used to collect the fluorescence in epifluorescence mode. A 488 nm long pass filter was used to eliminate any excitation wavelength, and a 100 μm pinhole to eliminate out-of-focus light. The fluorescence emission was split into two single photon sensitive avalanche photodiodes (MPD-SPAD) using a 50/50 beam splitting cube and the samples were analyzed under a 60X magnification, 1.2 numerical aperture (NA) objective (Olympus).

### Time-resolved fluorescence spectroscopy

Time-resolved fluorescence spectroscopic studies were carried out using the tryptophan variants of WT α-Syn (V3W, V71W, A124W, and A140W). 200 μM of each Trp variant in presence of 10 % PEG-8000 (pH 7.4) were incubated in a closed moist chamber at 37°C. 885 nm radiation from a Ti-sapphire laser (Mai Tai HP, Spectra Physics); pulsing at 80 MHz was used to irradiate the samples after frequency tripling to 295 nm. For the fluorescence lifetime experiments, the emission polarizer (Glan-Thompson) was set at a magic angle (54.7 °) with respect to the plane of the excitation beam to avoid polarization effects. For fluorescence anisotropy studies, however, it was kept at either parallel (0°) or perpendicular (90°) orientation with respect to the plane of excitation. The emission was recorded at 340 nm using a hybrid PMT detector (HPM-100-40, Becker and Hickl) operating at 83 % gain voltage. Single photon counting was initiated by a photodiode (APM-400, Becker and Hickl) and read by a TCSPC card (SPC-630, Becker and Hickl). Time per channel was 48.8 ps for our measurements. A colloidal mixture of a silica-based scattering solution (LUDOX) was used to collect the Instrument Response Function (IRF). The fluorescence lifetime of the aqueous solution of NATA (2.5-2.8 ns), which acts as a control for free Trp molecule in solution was collected routinely as a reference. The peak counts for all decay measurements were taken up to 10,000 for an acquisition window of 50 ns. The lifetime and fluorescence anisotropy measurements were taken both before and after LLPS, and analyzed to measure the mean lifetime and the rotational correlation time (Supplementary Information).

### Nuclear Magnetic Resonance (NMR) spectroscopy

Proteins were expressed in M9 minimal media with ^15^N labeled NH_4_Cl for NMR studies and purified as mentioned above. NMR experiments were recorded at 288 K on a 750 MHz NMR spectrophotometer equipped with a triple resonance 5 mm TXI probe (Bruker, USA). Two dimensional ^1^H-^15^N heteronuclear single quantum correlation spectroscopy (HSQC) of 200 μM α-Syn WT and A53T in 20 mM phosphate buffer containing 10 % PEG (pH 7.4) was recorded on the first five days (day 0, day 1, day 2, day 3, day 4) and day 20 of phase separation. Important to note that LLPS experiment was done in NMR tube only, where phase separation during the time was confirmed by analyzing small aliquot under the microscope. Each spectrum was acquired with 2046 and 256 data points in the direct and indirect dimensions. The spectra were recorded and processed using Bruker Topspin 3.5 software. The spectra were analyzed and the intensities of the amide cross-peaks obtained using CCP-NMR (Collaborative Computational Project for NMR).

### *In vitro* Förster resonance energy transfer (FRET) experiment

Trp (donor) and DTNB-labeled Cys (acceptor) pair was used to monitor time-dependent FRET. Briefly, 200 μM of tryptophan variants (V3W, V71W and A124W) were mixed with an equimolar concentration of respective DTNB-labeled V3C, V74C and A124C proteins in presence of 10 % PEG. During incubation, the samples were excited at 295 nm and the decrease in Trp fluorescence emission due to DTNB quenching was recorded using a spectrofluorimeter (FluoroMax-4, HORIBA Scientific, USA) in the range of 300 - 450 nm. Appropriate non-FRET pair controls (only donor, only acceptor) were taken for each experiment. The intermolecular FRET efficiency was calculated as the ratio of the fluorescence emission intensity of Trp in presence of DTNB (F_DA_) and Trp alone (F_D_);

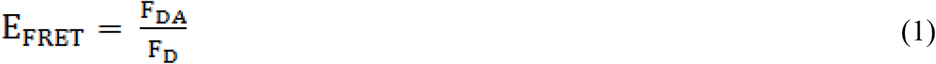

The distance (R) between the FRET donor (Trp) and acceptor (DTNB) can be calculated using the following equation;

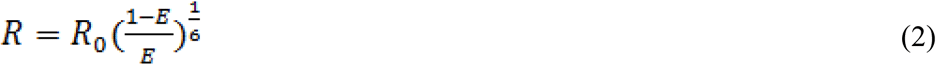

where R_0_ is the distance between Trp and DTNB in maximum denaturing conditions (in presence of 6 M Guanidine hydrochloride). The value of R_0_ is reported to be 23 Å (Bhatia et al., 2018).

For determining FRET in a single liquid droplet, spatially-resolved fluorescence imaging was employed. 1:1 Fluorescein and rhodamine maleimide (ThermoFisher Scientific, USA) labeled α-Syn containing single liquid droplet was imaged using a home-built epifluorescence total internal reflection fluorescence microscopy (Nikon Eclipse TE2000-U). 488 nm and 532 nm DPSS lasers (OXXIUS, model: ACX-CTRB and LASERGLOW, model: LRS-0532-PFM-00200-03, respectively) were used to illuminate a region of interest through a 1.49 NA, 60X TIRF objective lens (Nikon, Japan). The emission was collected and passed through the appropriate dichroic/long pass filters, and imaged through an air-cooled CCD camera (DVC 1412AM). The colocalization of fluorescein and rhodamine-labeled droplet was done by collecting the emission at two energetically separated detection channels, using 515 − 565 and 590 − 700 nm band-pass emission filters, respectively, while excited at 488 nm. Typically, images were collected at 50 ms exposure and averaged over 20 frames before further image processing. The background correction has been done with 50 pixels rolling ball radius. The spatially resolved spectroscopy of liquid droplets was done by the combination of a narrow slit and a transmission grating (70 g/mm) in the emission path before the sCMOS camera (ORCA-Flash4.0 V3, model: C13440-20CU). All the spectrally resolved images were collected at identical excitation power (5W/cm^2^) of 488 nm and 532 nm lasers at the same exposure time (0.3s). All measurements were performed at 295 K and the data were analyzed using ImageJ (NIH) and OriginPro 8 (OriginLab, USA) (Supplementary Information).

### α-Syn LLPS *in cells*

HeLa cells have been extensively used for studying LLPS in cells that over-express the protein of interest (Maharana et al., 2018; Maucuer et al., 2018; Molliex et al., 2015). Using a similar concept, we used HeLa cells stably expressing C4-α-Syn for our study. The details for the generation of the cell line are provided in Supplementary Information. The cells were cultured at 37°C in 5 % CO_2_ in DMEM (GIBCO) supplemented with 10 % FBS (GIBCO) and 20 ng of doxycycline hyclate (Sigma, USA). After 16 h of culturing, 10 mM of ferric ammonium citrate dissolved in water was used to induce stress. The cells were stained with FlAsH-EDT_2_ using the previously described method (Hoffmann et al., 2010). Briefly, cells were incubated with 1 μM FlAsH (Cayman Chemicals, USA) and 10 mM EDT in Opti-MEM (Invitrogen, USA), for 30 min at 37 °C and 5 % CO_2_. Subsequently, the unbound dye was removed by washing for 10 min with 100 mM EDT prepared in Opti-MEM. The images were captured using Zeiss Axio-Observer Z1 laser scanning confocal microscope (inverted) equipped with iPlan-Apochromat 63X/1.4 NA oil immersion objective. FlAsH was excited with an Ar-ion laser at 488 nm and detected with a 515 long-pass filter. Image processing and quantification were performed with Imaris 7.6.4 (BitPlane, Switzerland) and ImageJ (NIH). Annexin V-PI (BD Biosciences) apoptosis assay, MitoTracker Red (Catalog no M7513, Thermo Fisher Scientific, USA), and LysoTracker Red DND-99 (Catalog no. L7528, ThermoFisher Scientific) staining was performed as per the manufacturer’s protocol.

## SUPPLEMENTAL INFORMATION

Supplemental information includes Supplemental Experimental Procedures, Result, eight figures, and four movies.

## DECLARATION OF INTEREST

The authors declare no conflict of interests.

## AUTHOR CONTRIBUTIONS

S.R, R.K, L.G, K.P, J.M, R. Panigrahi, D.C, S.B, S.M. A.S., and S. Maiti performed the *in vitro* and *in silico* experiments. N.S, S.P, D.D, J.G. and A. N. performed the *in cell* experiments. All authors participated for analyzing data. The study was conceived by S.K.M and designed by S.K.M, G.K, R.P, A.K, A.C and R.R. S.K.M wrote the manuscript with the help of S.R., N.S., S.P. and S.M. All authors read and approved the manuscript.

## ACKNOWLEDGMENT

We thank Prof. Roop Malik (TIFR, Mumbai) and Prof. Madan Rao (NCBS, Bangalore) for critical inputs on the manuscript. We also acknowledge IIT Bombay central facilities for TEM, FACS, and Confocal microscopy for performing the experiments. We thank Rama Reddy and Sreemantee Sen (NCBS, Bangalore) for their help in TCSPC microscopy experiments. We acknowledge Prof. Charles Glabe, UC Urvine, USA for the kind gift of OC antibody. The authors acknowledge DBT [BT/PR22749/BRB/10/1576/2016], Government of India for financial support, and N.S acknowledges DST SERB [PDF/2016/003736] for funding.

## REFERENCES

Aguzzi, A., and Altmeyer, M. (2016). Phase Separation: Linking Cellular Compartmentalization to Disease. Trends Cell Biol. 26, 547–558.

Alberti, S. (2017). Phase separation in biology. Curr. Biol. 27, R1097–R1102.

Ambadipudi, S., Biernat, J., Riedel, D., Mandelkow, E., and Zweckstetter, M. (2017). Liquid-liquid phase separation of the microtubule-binding repeats of the Alzheimer-related protein Tau. Nat. Commun. 8, 275.

Banani, S.F., Lee, H.O., Hyman, A.A., and Rosen, M.K. (2017). Biomolecular condensates: organizers of cellular biochemistry. Nat. Rev. Mol. Cell Biol. 18, 285–298.

Banani, S.F., Rice, A.M., Peeples, W.B., Lin, Y., Jain, S., Parker, R., and Rosen, M.K. (2016). Compositional Control of Phase-Separated Cellular Bodies. Cell 166, 651–663.

Berry, J., Weber, S.C., Vaidya, N., Haataja, M., and Brangwynne, C.P. (2015). RNA transcription modulates phase transition-driven nuclear body assembly. Proc. Natl. Acad. Sci. U. S. A. 112, E5237–5245.

Bhatia, S., Krishnamoorthy, G., and Udgaonkar, J.B. (2018). Site-specific time-resolved FRET reveals local variations in the unfolding mechanism in an apparently two-state protein unfolding transition. Phys. Chem. Chem. Phys. 20, 3216–3232.

Boeynaems, S., Alberti, S., Fawzi, N.L., Mittag, T., Polymenidou, M., Rousseau, F., Schymkowitz, J., Shorter, J., Wolozin, B., Van Den Bosch, L., et al. (2018). Protein Phase Separation: A New Phase in Cell Biology. Trends Cell Biol. 28, 420–435.

Brangwynne, C.P., Eckmann, C.R., Courson, D.S., Rybarska, A., Hoege, C., Gharakhani, J., Julicher, F., and Hyman, A.A. (2009). Germline P granules are liquid droplets that localize by controlled dissolution/condensation. Science 324, 1729–1732.

Brangwynne, C.P., Mitchison, T.J., and Hyman, A.A. (2011). Active liquid-like behavior of nucleoli determines their size and shape in Xenopus laevis oocytes. Proc. Natl. Acad. Sci. U. S. A. 108, 4334–4339.

Breydo, L., Wu, J.W., and Uversky, V.N. (2012). α-synuclein misfolding and Parkinson’s disease. Biochim. Biophys. Acta 1822, 261–285.

Buell, A.K., Galvagnion, C., Gaspar, R., Sparr, E., Vendruscolo, M., Knowles, T.P., Linse, S., and Dobson, C.M. (2014). Solution conditions determine the relative importance of nucleation and growth processes in alpha-synuclein aggregation. Proc. Natl. Acad. Sci. U. S. A. 111, 7671–7676.

Burke, K.A., Janke, A.M., Rhine, C.L., and Fawzi, N.L. (2015). Residue-by-Residue View of In Vitro FUS Granules that Bind the C-Terminal Domain of RNA Polymerase II. Mol. Cell 60, 231–241.

Conicella, A.E., Zerze, G.H., Mittal, J., and Fawzi, N.L. (2016). ALS Mutations Disrupt Phase Separation Mediated by alpha-Helical Structure in the TDP-43 Low-Complexity C-Terminal Domain. Structure 24, 1537–1549.

Conway, K.A., Harper, J.D., and Lansbury, P.T. (1998). Accelerated in vitro fibril formation by a mutant alpha-synuclein linked to early-onset Parkinson disease. Nat. Medicine 4, 1318–1320.

Conway, K.A., Lee, S.J., Rochet, J.C., Ding, T.T., Williamson, R.E., and Lansbury, P.T., Jr. (2000). Acceleration of oligomerization, not fibrillization, is a shared property of both α-synuclein mutations linked to early-onset Parkinson’s disease: implications for pathogenesis and therapy. Proc. Natl. Acad. Sci. U. S. A. 97, 571–576.

Conway, K.A., Rochet, J.C., Bieganski, R.M., and Lansbury, P.T., Jr. (2001). Kinetic stabilization of the α-synuclein protofibril by a dopamine-α-synuclein adduct. Science 294, 1346–1349.

Cookson, M.R. (2005). The biochemistry of Parkinson’s disease. Annu. Rev. Biochem. 74, 29–52.

Dedmon, M.M., Lindorff-Larsen, K., Christodoulou, J., Vendruscolo, M., and Dobson, C.M. (2005). Mapping long-range interactions in alpha-synuclein using spin-label NMR and ensemble molecular dynamics simulations. J. Am. Chem. Soc. 127, 476–477.

Elbaum-Garfinkle, S., Kim, Y., Szczepaniak, K., Chen, C.C., Eckmann, C.R., Myong, S., and Brangwynne, C.P. (2015). The disordered P granule protein LAF-1 drives phase separation into droplets with tunable viscosity and dynamics. Proc. Natl. Acad. Sci. U. S. A. 112, 7189–7194.

Eliezer, D., Kutluay, E., Bussell, R., Jr., and Browne, G. (2001). Conformational properties of alpha-synuclein in its free and lipid-associated states. J. Mol. Biol. 307, 1061–1073.

Falahati, H., and Wieschaus, E. (2017). Independent active and thermodynamic processes govern the nucleolus assembly in vivo. Proc. Natl. Acad. Sci. U. S. A. 114, 1335–1340.

Feric, M., and Brangwynne, C.P. (2013). A nuclear F-actin scaffold stabilizes ribonucleoprotein droplets against gravity in large cells. Nat. Cell Biol. 15, 1253–1259.

Feric, M., Vaidya, N., Harmon, T.S., Mitrea, D.M., Zhu, L., Richardson, T.M., Kriwacki, R.W., Pappu, R.V., and Brangwynne, C.P. (2016). Coexisting Liquid Phases Underlie Nucleolar Subcompartments. Cell 165, 1686–1697.

Fonin, A.V., Darling, A.L., Kuznetsova, I.M., Turoverov, K.K., and Uversky, V.N. (2018). Intrinsically disordered proteins in crowded milieu: when chaos prevails within the cellular gumbo. Cell. Mol. life Sci. 75, 3907–3929.

Fusco, D., Accornero, N., Lavoie, B., Shenoy, S.M., Blanchard, J.M., Singer, R.H., and Bertrand, E. (2003). Single mRNA molecules demonstrate probabilistic movement in living mammalian cells. Curr. Biol. 13, 161–167.

Garcia-Mata, R., Bebok, Z., Sorscher, E.J., and Sztul, E.S. (1999). Characterization and dynamics of aggresome formation by a cytosolic GFP-chimera. J. Cell Biol. 146, 1239–1254.

Garcia-Mata, R., Capdevielle, J., Guillemot, J.C., Ferrara, P., Conde, R.D., and Sanllorenti, P.M. (1997). Protein depletion and refeeding change the proportion of mouse liver glutathione S-transferase subunits. Biochim. Biophys. Acta 1357, 272–280.

Ghosh, D., Sahay, S., Ranjan, P., Salot, S., Mohite, G.M., Singh, P.K., Dwivedi, S., Carvalho, E., Banerjee, R., Kumar, A., et al. (2014). The newly discovered Parkinson’s disease associated Finnish mutation (A53E) attenuates α-synuclein aggregation and membrane binding. Biochemistry 53, 6419–6421.

Giasson, B.I., Murray, I.V., Trojanowski, J.Q., and Lee, V.M. (2001). A hydrophobic stretch of 12 amino acid residues in the middle of α-synuclein is essential for filament assembly. J. Biol. Chem. 276, 2380–2386.

Goedert, M. (2001). α-synuclein and neurodegenerative diseases. Nat. Rev. Neurosci. 2, 492–501.

Greenbaum, E.A., Graves, C.L., Mishizen-Eberz, A.J., Lupoli, M.A., Lynch, D.R., Englander, S.W., Axelsen, P.H., and Giasson, B.I. (2005). The E46K mutation in α-synuclein increases amyloid fibril formation. J. Biol. Chem. 280, 7800–7807.

Greenspan, P., Mayer, E.P., and Fowler, S.D. (1985). Nile red: a selective fluorescent stain for intracellular lipid droplets. J. Cell Biol. 100, 965–973.

Guerrero-Ferreira, R., Taylor, N.M., Mona, D., Ringler, P., Lauer, M.E., Riek, R., Britschgi, M., and Stahlberg, H. (2018). Cryo-EM structure of α-synuclein fibrils. eLife 7, e36402.

Han, T.W., Kato, M., Xie, S., Wu, L.C., Mirzaei, H., Pei, J., Chen, M., Xie, Y., Allen, J., Xiao, G., et al. (2012). Cell-free formation of RNA granules: bound RNAs identify features and components of cellular assemblies. Cell 149, 768–779.

Hoffmann, C., Gaietta, G., Zurn, A., Adams, S.R., Terrillon, S., Ellisman, M.H., Tsien, R.Y., and Lohse, M.J. (2010). Fluorescent labeling of tetracysteine-tagged proteins in intact cells. Nat. protocols 5, 1666–1677.

Hughes, M.P., Sawaya, M.R., Boyer, D.R., Goldschmidt, L., Rodriguez, J.A., Cascio, D., Chong, L., Gonen, T., and Eisenberg, D.S. (2018). Atomic structures of low-complexity protein segments reveal kinked beta sheets that assemble networks. Science 359, 698–701.

Hyman, A.A., Weber, C.A., and Julicher, F. (2014). Liquid-liquid phase separation in biology. Annu. Rev. Cell Dev. Biol. 30, 39–58.

Irtegun, S., Ramdzan, Y.M., Mulhern, T.D., and Hatters, D.M. (2011). ReAsH/FlAsH labeling and image analysis of tetracysteine sensor proteins in cells. J. Vis Exp. 31, e2857.

Jha, N.N., Kumar, R., Panigrahi, R., Navalkar, A., Ghosh, D., Sahay, S., Mondal, M., Kumar, A., and Maji, S.K. (2017). Comparison of alpha-Synuclein Fibril Inhibition by Four Different Amyloid Inhibitors. ACS Chem Neurosci. 8, 2722–2733.

Jha, S.K., and Udgaonkar, J.B. (2009). Direct evidence for a dry molten globule intermediate during the unfolding of a small protein. Proc. Natl. Acad. Sci. U. S. A. 106, 12289–12294.

Johnston, J.A., Illing, M.E., and Kopito, R.R. (2002). Cytoplasmic dynein/dynactin mediates the assembly of aggresomes. Cell Motil. Cytoskeleton 53, 26–38.

Kaiser, T.E., Intine, R.V., and Dundr, M. (2008). De Novo Formation of a Subnuclear Body. Science 322, 1713–1717.

Kaizuka, Y., Douglass, A.D., Varma, R., Dustin, M.L., and Vale, R.D. (2007). Mechanisms for segregating T cell receptor and adhesion molecules during immunological synapse formation in Jurkat T cells. Proc. Natl. Acad. Sci. U. S. A. 104, 20296–20301.

Kato, M., Han, T.W., Xie, S., Shi, K., Du, X., Wu, L.C., Mirzaei, H., Goldsmith, E.J., Longgood, J., Pei, J., et al. (2012). Cell-free formation of RNA granules: low complexity sequence domains form dynamic fibers within hydrogels. Cell 149, 753–767.

Kayed, R., Head, E., Sarsoza, F., Saing, T., Cotman, C.W., Necula, M., Margol, L., Wu, J., Breydo, L., Thompson, J.L., et al. (2007). Fibril specific, conformation dependent antibodies recognize a generic epitope common to amyloid fibrils and fibrillar oligomers that is absent in prefibrillar oligomers. Mol. Neurodegener. 2, 18.

Kruger, R., Kuhn, W., Muller, T., Woitalla, D., Graeber, M., Kosel, S., Przuntek, H., Epplen, J.T., Schols, L., and Riess, O. (1998). Ala30Pro mutation in the gene encoding alpha-synuclein in Parkinson’s disease. Nat. Genetics 18, 106–108.

Kumar, R., Das, S., Mohite, G.M., Rout, S.K., Halder, S., Jha, N.N., Ray, S., Mehra, S., Agarwal, V., and Maji, S.K. (2018). Cytotoxic Oligomers and Fibrils Trapped in a Gel-like State of alpha-Synuclein Assemblies. Angew. Chem. Intl. Ed. 57, 5262–5266.

Kwon, I., Kato, M., Xiang, S., Wu, L., Theodoropoulos, P., Mirzaei, H., Han, T., Xie, S., Corden, J.L., and McKnight, S.L. (2013). Phosphorylation-regulated binding of RNA polymerase II to fibrous polymers of low-complexity domains. Cell 155, 1049–1060.

Lee, H.J., Choi, C., and Lee, S.J. (2002). Membrane-bound α-synuclein has a high aggregation propensity and the ability to seed the aggregation of the cytosolic form. J. Biol. Chem. 277, 671–678.

Letunic, I., and Bork, P. (2018). 20 years of the SMART protein domain annotation resource. Nucleic Acids Res. 46, D493–D496.

Li, B.S., Ge, P., Murray, K.A., Sheth, P., Zhang, M., Nair, G., Sawaya, M.R., Shin, W.S., Boyer, D.R., Ye, S.L., et al. (2018a). Cryo-EM of full-length α-synuclein reveals fibril polymorphs with a common structural kernel. Nat. Commun. 9, 3609.

Li, H.R., Chiang, W.C., Chou, P.C., Wang, W.J., and Huang, J.R. (2018b). TAR DNA-binding protein 43 (TDP-43) liquid-liquid phase separation is mediated by just a few aromatic residues. J. Biol. Chem. 293, 6090–6098.

Li, P., Banjade, S., Cheng, H.C., Kim, S., Chen, B., Guo, L., Llaguno, M., Hollingsworth, J.V., King, D.S., Banani, S.F., et al. (2012). Phase transitions in the assembly of multivalent signalling proteins. Nature 483, 336–340.

Lin, Y., Protter, D.S., Rosen, M.K., and Parker, R. (2015). Formation and Maturation of Phase-Separated Liquid Droplets by RNA-Binding Proteins. Mol. Cell 60, 208–219.

Maharana, S., Wang, J., Papadopoulos, D.K., Richter, D., Pozniakovsky, A., Poser, I., Bickle, M., Rizk, S., Guillen-Boixet, J., Franzmann, T.M., et al. (2018). RNA buffers the phase separation behavior of prion-like RNA binding proteins. Science 360, 918–921.

Matsuzaki, M., Hasegawa, T., Takeda, A., Kikuchi, A., Furukawa, K., Kato, Y., and Itoyama, Y. (2004). Histochemical features of stress-induced aggregates in α-synuclein overexpressing cells. Brain Res. 1004, 83–90.

Maucuer, A., Desforges, B., Joshi, V., Boca, M., Kretov, D.A., Hamon, L., Bouhss, A., Curmi, P.A., and Pastre, D. (2018). Microtubules as platforms for probing liquid-liquid phase separation in cells - application to RNA-binding proteins. J. Cell Sci. 131, 214692.

Meszaros, B., Erdos, G., and Dosztanyi, Z. (2018). IUPred2A: context-dependent prediction of protein disorder as a function of redox state and protein binding. Nucleic Acids Res 46, W329–W337.

Mitrea, D.M., Cika, J.A., Guy, C.S., Ban, D., Banerjee, P.R., Stanley, C.B., Nourse, A., Deniz, A.A., and Kriwacki, R.W. (2016). Nucleophosmin integrates within the nucleolus via multi-modal interactions with proteins displaying R-rich linear motifs and rRNA. eLife 5, e13571.

Mitrea, D.M., and Kriwacki, R.W. (2016). Phase separation in biology; functional organization of a higher order. Cell Commun. Signal. 14, 1.

Molliex, A., Temirov, J., Lee, J., Coughlin, M., Kanagaraj, A.P., Kim, H.J., Mittag, T., and Taylor, J.P. (2015). Phase separation by low complexity domains promotes stress granule assembly and drives pathological fibrillization. Cell 163, 123–133.

Monahan, Z., Ryan, V.H., Janke, A.M., Burke, K.A., Rhoads, S.N., Zerze, G.H., O’Meally, R., Dignon, G.L., Conicella, A.E., Zheng, W., et al. (2017). Phosphorylation of the FUS low-complexity domain disrupts phase separation, aggregation, and toxicity. EMBO J. 36, 2951–2967.

Munder, M.C., Midtvedt, D., Franzmann, T., Nuske, E., Otto, O., Herbig, M., Ulbricht, E., Muller, P., Taubenberger, A., Maharana, S., et al. (2016). A pH-driven transition of the cytoplasm from a fluid-to a solid-like state promotes entry into dormancy. eLife 5, e09347.

Murakami, T., Qamar, S., Lin, J.Q., Schierle, G.S., Rees, E., Miyashita, A., Costa, A.R., Dodd, R.B., Chan, F.T., Michel, C.H., et al. (2015). ALS/FTD Mutation-Induced Phase Transition of FUS Liquid Droplets and Reversible Hydrogels into Irreversible Hydrogels Impairs RNP Granule Function. Neuron 88, 678–690.

Narhi, L., Wood, S.J., Steavenson, S., Jiang, Y., Wu, G.M., Anafi, D., Kaufman, S.A., Martin, F., Sitney, K., Denis, P., et al. (1999). Both familial Parkinson’s disease mutations accelerate alpha-synuclein aggregation. J. Biol. Chem. 274, 9843–9846.

Nott, T.J., Petsalaki, E., Farber, P., Jervis, D., Fussner, E., Plochowietz, A., Craggs, T.D., Bazett-Jones, D.P., Pawson, T., Forman-Kay, J.D., et al. (2015). Phase transition of a disordered nuage protein generates environmentally responsive membraneless organelles. Mol. Cell 57, 936–947.

Ostrerova-Golts, N., Petrucelli, L., Hardy, J., Lee, J.M., Farer, M., and Wolozin, B. (2000). The A53T alpha-synuclein mutation increases iron-dependent aggregation and toxicity. J. Neurosci. 20, 6048–6054.

Pak, C.W., Kosno, M., Holehouse, A.S., Padrick, S.B., Mittal, A., Ali, R., Yunus, A.A., Liu, D.R., Pappu, R.V., and Rosen, M.K. (2016). Sequence Determinants of Intracellular Phase Separation by Complex Coacervation of a Disordered Protein. Mol. Cell 63, 72–85.

Pasanen, P., Myllykangas, L., Siitonen, M., Raunio, A., Kaakkola, S., Lyytinen, J., Tienari, P.J., Poyhonen, M., and Paetau, A. (2014). Novel alpha-synuclein mutation A53E associated with atypical multiple system atrophy and Parkinson’s disease-type pathology. Neurobiol. Aging 35, 2180 e2181–2185.

Patel, A., Lee, H.O., Jawerth, L., Maharana, S., Jahnel, M., Hein, M.Y., Stoynov, S., Mahamid, J., Saha, S., Franzmann, T.M., et al. (2015). A Liquid-to-Solid Phase Transition of the ALS Protein FUS Accelerated by Disease Mutation. Cell 162, 1066–1077.

Paxinou, E., Chen, Q., Weisse, M., Giasson, B.I., Norris, E.H., Rueter, S.M., Trojanowski, J.Q., Lee, V.M., and Ischiropoulos, H. (2001). Induction of α-synuclein aggregation by intracellular nitrative insult. J. Neurosci. 21, 8053–8061.

Polymeropoulos, M.H., Lavedan, C., Leroy, E., Ide, S.E., Dehejia, A., Dutra, A., Pike, B., Root, H., Rubenstein, J., Boyer, R., et al. (1997). Mutation in the α-synuclein gene identified in families with Parkinson’s disease. Science 276, 2045–2047.

Reichheld, S.E., Muiznieks, L.D., Keeley, F.W., and Sharpe, S. (2017). Direct observation of structure and dynamics during phase separation of an elastomeric protein. Proc. Natl. Acad. Sci. U. S. A. 114, E4408–E4415.

Riback, J.A., Katanski, C.D., Kear-Scott, J.L., Pilipenko, E.V., Rojek, A.E., Sosnick, T.R., and Drummond, D.A. (2017). Stress-Triggered Phase Separation Is an Adaptive, Evolutionarily Tuned Response. Cell 168, 1028–1040 e1019.

Roberti, M.J., Bertoncini, C.W., Klement, R., Jares-Erijman, E.A., and Jovin, T.M. (2007). Fluorescence imaging of amyloid formation in living cells by a functional, tetracysteine-tagged α-synuclein. Nat. Methods 4, 345–351.

Sahay, S., Ghosh, D., Dwivedi, S., Anoop, A., Mohite, G.M., Kombrabail, M., Krishnamoorthy, G., and Maji, S.K. (2015). Familial Parkinson disease-associated mutations alter the site-specific microenvironment and dynamics of α-synuclein. J. Biol. Chem. 290, 7804–7822.

Sahay, S., Krishnamoorthy, G., and Maji, S.K. (2016). Site-specific structural dynamics of α-Synuclein revealed by time-resolved fluorescence spectroscopy: a review. Meth. Appl. Fluoresc. 4, 042002.

Schuster, B.S., Reed, E.H., Parthasarathy, R., Jahnke, C.N., Caldwell, R.M., Bermudez, J.G., Ramage, H., Good, M.C., and Hammer, D.A. (2018). Controllable protein phase separation and modular recruitment to form responsive membraneless organelles. Nat. Commun. 9, 2985.

Shin, Y., Berry, J., Pannucci, N., Haataja, M.P., Toettcher, J.E., and Brangwynne, C.P. (2017). Spatiotemporal control of intracellular phase transitions using light-activated optoDroplets. Cell 168, 159–171 e114.

Shin, Y., and Brangwynne, C.P. (2017). Liquid phase condensation in cell physiology and disease. Science 357.

Shtilerman, M.D., Ding, T.T., and Lansbury, P.T., Jr. (2002). Molecular crowding accelerates fibrillization of α-synuclein: could an increase in the cytoplasmic protein concentration induce Parkinson’s disease? Biochemistry 41, 3855–3860.

Singh, P.K., Kotia, V., Ghosh, D., Mohite, G.M., Kumar, A., and Maji, S.K. (2013). Curcumin modulates α-synuclein aggregation and toxicity. ACS Chem. Neurosci. 4, 393–407.

Spillantini, M.G., Schmidt, M.L., Lee, V.M., Trojanowski, J.Q., Jakes, R., and Goedert, M. (1997). α-synuclein in Lewy bodies. Nature 388, 839–840.

Tarantino, N., Tinevez, J.Y., Crowell, E.F., Boisson, B., Henriques, R., Mhlanga, M., Agou, F., Israel, A., and Laplantine, E. (2014). TNF and IL-1 exhibit distinct ubiquitin requirements for inducing NEMO-IKK supramolecular structures. J. Cell Biol. 204, 231–245.

Uversky, V.N., Li, J., and Fink, A.L. (2001). Evidence for a partially folded intermediate in alpha-synuclein fibril formation. J. Biol. Chem. 276, 10737–10744.

Uversky, V.N., Li, J., Souillac, P., Millett, I.S., Doniach, S., Jakes, R., Goedert, M., and Fink, A.L. (2002). Biophysical properties of the synucleins and their propensities to fibrillate: inhibition of alpha-synuclein assembly by β- and Υ-synucleins. J. Biol. Chem. 277, 11970–11978.

Vilar, M., Chou, H.T., Luhrs, T., Maji, S.K., Riek-Loher, D., Verel, R., Manning, G., Stahlberg, H., and Riek, R. (2008). The fold of alpha-synuclein fibrils. Proc. Natl. Acad. Sci. U. S. A. 105, 8637–8642.

Volles, M.J., and Lansbury, P.T. (2003). Zeroing in on the pathogenic form of α-synuclein and its mechanism of neurotoxicity in Parkinson’s disease. Biochemistry 42, 7871–7878.

Volles, M.J. and Lansbury, P.T. (2007) Relationships between the sequence of alpha-synuclein and its membrane affinity, fibrillization propensity, and yeast toxicity. J. Mol. Biol. 366, 1510–1522.

Voorhees, P.W. (1992). Ostwald Ripening of 2-Phase Mixtures. Annu. Rev. Mater. Sci. 22, 197–215.

Wang, M., Tao, X., Jacob, M.D., Bennett, C.A., Ho, J.J.D., Gonzalgo, M.L., Audas, T.E., and Lee, S. (2018). Stress-Induced Low Complexity RNA Activates Physiological Amyloidogenesis. Cell Rep. 24, 1713–1721 e1714.

Wegmann, S., Eftekharzadeh, B., Tepper, K., Zoltowska, K.M., Bennett, R.E., Dujardin, S., Laskowski, P.R., MacKenzie, D., Kamath, T., Commins, C., et al. (2018). Tau protein liquid-liquid phase separation can initiate tau aggregation. EMBO J. 37, e98049.

Whitt, M.A., and Mire, C.E. (2011). Utilization of fluorescently-labeled tetracysteine-tagged proteins to study virus entry by live cell microscopy. Methods 55, 127–136.

Winner, B., Jappelli, R., Maji, S.K., Desplats, P.A., Boyer, L., Aigner, S., Hetzer, C., Loher, T., Vilar, M., Campioni, S., et al. (2011). In vivo demonstration that α-synuclein oligomers are toxic. Proc. Natl. Acad. Sci. U. S. A. 108, 4194–4199.

Wu, K.P., Weinstock, D.S., Narayanan, C., Levy, R.M., and Baum, J. (2009). Structural reorganization of α-synuclein at low pH observed by NMR and REMD simulations. J. Mol. Biol. 391, 784–796.

Xiang, S., Kato, M., Wu, L.C., Lin, Y., Ding, M., Zhang, Y., Yu, Y., and McKnight, S.L. (2015). The LC domain of hnRNPA2 adopts similar conformations in hydrogel polymers, liquid-like droplets, and nuclei. Cell 163, 829–839.

Yan, C., and Pochan, D.J. (2010). Rheological properties of peptide-based hydrogels for biomedical and other applications. Chem. Soc. Rev. 39, 3528–3540.

Yasuda, T., Nakata, Y., and Mochizuki, H. (2013). α-Synuclein and neuronal cell death. Mol. Neurobiol. 47, 466–483.

Yi, J., Wu, X.S., Crites, T., and Hammer, J.A., 3rd (2012). Actin retrograde flow and actomyosin II arc contraction drive receptor cluster dynamics at the immunological synapse in Jurkat T cells. Mol. Biol. Cell 23, 834–852.

Zarranz, J.J., Alegre, J., Gomez-Esteban, J.C., Lezcano, E., Ros, R., Ampuero, I., Vidal, L., Hoenicka, J., Rodriguez, O., Atares, B., et al. (2004). The new mutation, E46K, of α-synuclein causes Parkinson and Lewy body dementia. Annals Neurol. 55, 164–173.

Zhang, H., Elbaum-Garfinkle, S., Langdon, E.M., Taylor, N., Occhipinti, P., Bridges, A.A., Brangwynne, C.P., and Gladfelter, A.S. (2015). RNA Controls PolyQ Protein Phase Transitions. Mol. Cell 60, 220–230.

