## Supplementary Information for "Liquid-liquid phase separation and liquid-to-solid transition mediate α-synuclein amyloid fibril containing hydrogel formation"

### Materials and Methods:

#### Reagents

All the chemicals and reagents were of the analytical grade from Sigma (USA) and HiMedia (India), unless otherwise specified. Fluorescein, rhodamine dyes and DTNB were obtained from ThermoFisher Scientific. FlAsH-EDT$_2$ (CAS No. 212118-77-9) was purchased from Cayman Chemicals (USA). Cell culture media, fetal bovine serum were obtained from GIBCO, USA. The following antibodies were used in the study: Anti-FLAG M2 antibody (catalog no. F1804, Sigma), α-tubulin, clone DM1A (catalog no. 05-829, Merck, USA). The HRP-tagged secondary antibody and fluorophore-labeled secondary antibodies were purchased from Calbiochem and Invitrogen (USA), respectively. Amyloid fibril-specific OC antibody was a kind gift from Prof. Charles G. Glabe (University of California, Irvine, CA). Dopamine hydrochloride, rotenone, 3-(4, 5-dimethylthiazol-2-yl)-2, 5-diphenyltetrazolium bromide (MTT), PEG-8000, doxycycline hyclate, ammonium ferric citrate were procured from Sigma (USA). The kits include: MitoTracker Red CM-H2XROS (Catalog no M7513, Thermo Fisher Scientific, USA), ProteoStat-aggresome detection kit (Enzo Life sciences, USA), LysoTracker Red DND-99 (Catalog no. L7528, ThermoFisher Scientific, USA), FITC Annexin V apoptosis detection kit (Catalog no. 556547, BD Biosciences, USA).

#### Protein expression and purification

WT α-Syn and its variants (A53T, core α-Syn (30-110), Trp mutants (3W, 71W, 124W, 140W), Cys mutants (3C, 74C, 124C), S129E phosphomimetic), β- and γ-Syn were expressed and purified using the standard protocols as described earlier (Volles et al., 2007; Singh et al., 2013). Briefly, *E.coli* BL21 (DE3) competent cells were transformed with the cloned plasmids and expression was induced by the addition of 1 mM IPTG. Subsequently, the cells were centrifuged and lysed using a probe sonicator (Sonics and Materials Inc., USA) and heat denatured for 20 minutes. For the Cys variants, sonication and subsequent steps were performed in the presence of 1 mM DTT in a reducing environment. From the supernatant, DNA was precipitated using 10% (w/v) streptomycin sulfate and glacial acetic acid followed by saturated ammonium sulfate precipitation. Thereafter, the protein was washed thrice by 100 mM ammonium acetate and was precipitated from the ammonium acetate solution again using ethanol. The final protein pellet was dissolved in a minimum volume of 100 mM ammonium acetate and subsequently lyophilized. The $N^{15}$ labeled α-Syn proteins for the NMR studies were expressed and purified in M9 media with $N^{15}$ labeled $NH_4Cl$ as the sole nitrogen source.

#### Low Molecular Weight (LMW) protein preparation

Lyophilized protein was dissolved in 20 mM phosphate buffer (pH 7.4). The protein was solubilized by addition of 0.2 N NaOH and the final pH was adjusted to 7.4 using 2 M HCl with the help of micro-pH meter probe (Mettler Toledo, USA). The protein solution was centrifuged at 13,000 x g for 30 min at 4 $^{o}$C to remove insoluble aggregates. Thereafter, the supernatant was dialyzed overnight against the same buffer using 10 kDa cut off membranes (Sigma, USA) and subsequently passed through a pre-washed 100 kDa cutoff filter (Merck Millipore, USA) to remove any high-order aggregates. The flow-through i.e. the low molecular weight (LMW) solution (Singh et al., 2013) was collected, and the concentration was estimated by measuring the absorbance at 280 nm. The molar extinction coefficients (ε) used for calculating concentration of Trp mutants were 11400 $M^{-1}cm^{-1}$; for WT, A53T and Cys mutants, 5960 $M^{-1}cm^{-1}$; for γ- Syn, β-Syn and core Syn 1490 $M^{-1}cm^{-1}$, 5960 and 1490 $M^{-1}cm^{-1}$, respectively (ExPASy ProtParam, SIB).

**Light scattering measurement**

200 μM LMW protein of WT and A53T α-Syn along with 10% PEG (LLPS relevant conditions) were incubated at 37 °C in 2 ml microcentrifuge tubes (Eppendorf, USA) and used for light scattering measurements with excitation and emission at 350 nm. The measurements were carried out using a spectrofluorimeter (JASCO FP 8500, USA) with 2.5 nm excitation and emission slit width. The scattering was recorded for 30 seconds with an interval of 1 second. The magnitude of the scattering at $15^{th}$ second was plotted with respect to time. The experiments were performed with two independent sets.

**Protein labeling**

Fluorescein, rhodamine and DTNB labeling were done as per the manufacturer's instructions (ThermoFisher Scientific, USA). In brief, 10X molar excess of fluorescein/ rhodamine or DTNB (dissolved in DMSO) was added to the LMW protein. For FITC, the mixture was incubated for 4 h at room temperature (RT) followed by 6 h at 4°C with slow mixing with the help of a magnetic stirrer. For fluorescein-5-maleimide, rhodamine-C2-maleimide, NHS-rhodamine and DTNB, the protein was mixed with the dye at RT for ≥2 h. The excess dye was removed by dialysis against 20 mM phosphate buffer; pH 7.4 at 4°C for 48 h, with change of buffers every 5 h. For further experiments, we used 1:10 v/v ratio of labeled versus unlabeled protein, unless mentioned otherwise.

**Fluorescence and confocal microscopy**

The liquid-liquid phase separation and liquid droplet formation by α-Syn *in vitro* were visualized using a DMi8 microscope (Leica Microsystems, Germany) under DIC and fluorescence mode. The fluorophore-labeled droplets and ThioS binding were observed using appropriate fluorescence channels (488 nm for FITC, Fluorescein-5-maleimide and ThioS; 560 nm for rhodamine). Appropriate buffer controls for each experiment were kept for baseline fluorescence settings. The FRAP studies were performed using laser scanning confocal microscope (Zeiss Axio-Observer Z1 microscope (inverted)) equipped with iPlan-Apochromat 63X/1.4 NA oil immersion objective.

**Preparation of liposomes (SUVs)**

1-palmitoyl-2-oleoyl-sn-glycero-3-phosphate (POP) and 1-palmitoyl-2-oleoyl-sn-glycero-3-phosphocholine (POPC) (Avanti Polar Lipids Inc. USA) were mixed in a molar ratio of 1:3 (POP: POPC) and dissolved in chloroform. The solvent was evaporated by vacuum using a rotavap (Heidolph, Germany) with constant rotation of 50 rpm so that after evaporation of the chloroform; a thin lipid film is generated around the surface of the round bottom flask. The film was dissolved in 20 mM sodium-phosphate buffer (pH 7.4) by rotating the sample at 25 °C for ≥4 h. The final concentration of the lipid suspension was adjusted to 6 mM. The suspension was sonicated with a probe sonicator (Sonics and Materials Inc.; USA) at 40% amplitude for ~20 min to convert the large multilamellar vesicles (LMVs) into small unilamellar vesicles (SUVs). The average size of the vesicles was ~100 nm as confirmed by scanning electron microscopy and DLS studies (data not shown).

**Aggregation studies of various synuclein**

200 μM LMW-α-Syn dissolved in 20 mM phosphate buffer (pH 7.4) in presence of 10% PEG (standard LLPS condition) was incubated at 37 °C under static condition. At different time points, the solution was monitored for aggregation using Thioflavin T (ThT) to check for amyloid formation. Similar study was performed for A53T α-Syn. Important to note, the aggregation studies using ThT for β-, γ- and core-Syn, and α-Syn in the presence of various PD-associated factors were performed under rotating conditions (10 rpm) to facilitate the aggregation process. The LMW concentration used for β-, γ- and core-Syn was 200 μM.

**Thioflavin T (ThT) binding assay**

To monitor the α-Syn aggregation kinetics, 4 μl of 1 mM ThT was added to 20 μl protein solution (200 μM protein) used both for various aggregation as well as LLPS conditions, which was further diluted to 400 μl using 20 mM sodium phosphate buffer (pH 7.4). Using a spectrofluorimeter (JASCO FP 8500), the assay was performed at an excitation wavelength of 450 nm and the emission spectrum was collected in the range of 460 - 500 nm. A plot of emission maxima at 480 nm against time resulted in a sigmoidal curve.

**Thioflavin S (ThioS) staining**

10 μl of 0.0625% of ThioS (w/v) prepared in 20 mM sodium-phosphate buffer (pH 7.4) was mixed with 100 μl of LLPS reaction mixture (200 μM protein and 10% PEG-8000) and incubated at 37°C. At different time intervals, ThioS fluorescence was visualized under the DMi8 microscope (Leica Microsystems, Germany).

**Electron microscopy**

50 μM of protein solution was spotted on the carbon coated formvar grid (Electron Microscopy Sciences, USA) and incubated for 5 minutes. The grid was washed with Milli-Q water for 2 times and then stained with 5% aqueous uranyl formate solution and air dried before imaging.

For LLPS samples, the coverslip was removed from the slide and sample was transferred on the EM grid (Electron Microscopy Sciences, USA) directly. The grids were stained with uranyl formate without any Milli-Q wash and air-dried before imaging. The grids were subjected to transmission electron microscopy (TEM) imaging (Phillips CM-200, Amsterdam, Netherlands) at 200 kV and 6600X magnification. Images were recorded using Keen View Soft imaging system (Olympus, Tokyo, Japan). For scanning electron microscope (SEM) imaging, the LLPS and/or hydrogel samples were directly spotted onto a carbon coated grid directly and air dried and subjected to imaging (JSM-7600F, JEOL, JAPAN).

**Circular Dichroism (CD) spectroscopy**

10 µl of protein sample was diluted to 200 µl with 20 mM phosphate buffer, pH 7.4. The solution was transferred to a CD microcuvette (Hellma, Forest Hills, NY) of 0.1 cm path length and spectra was recorded from 260 to 200 nm at 25 °C with a speed of 100 nm/min with the help of a CD spectrophotometer (JASCO-1500). Appropriate controls were taken for each experiment. Each data was obtained in duplicates and three accumulations were taken for every sample. Buffer subtraction and smoothing of the raw data was done as per the manufacturer's instructions.

**Isolation of α-Syn components during LLPS and aggregation**

200 µM of α-Syn in 20 mM sodium phosphate buffer, pH 7.4, 0.02% sodium azide, in presence of 10% PEG was incubated in a 2 ml microcentrifuge tube (Eppendorf, USA) at 37°C without agitation for 30 days. At an interval of 5 days, aliquots were taken and checked for ThT fluorescence, liquid droplet formation (under phase contrast microscope) and CD spectroscopy. Rest of the aliquot was used for isolating α-Syn aggregated species according to our previously published protocol (Kumar et al., 2018). Briefly, samples were centrifuged at 100,000 g for 60 min (Beckman Coulter Optima MAX-XP Ultracentrifuge). Any fibril and higher order aggregates are retained in the pellet, whereas supernatant majorly constitutes oligomer and LMW fraction. The protein concentration in the supernatant was then determined by UV spectroscopy (Jasco V650) using molar absorptivity (ε) of 5960 $M^{-1}cm^{-1}$ for α-Syn, and the concentration of protein in pellet was back calculated by subtracting the protein concentration in supernatant from initial protein concentration (200 µM). Supernatant was further loaded onto pre-equilibrated 100 kDa cut-off filter (0.5 ml, Amicon Ultra, Merck Millipore) and centrifuged at 10,000 g for 30 min. The flow-through contains LMW form of α-Syn, while the oligomer is retained within the filter. The protein concentration in the flow-through was determined by UV spectroscopy (Jasco V650), and the oligomer concentration was obtained by subtracting LMW concentration from the supernatant. The individual components were then analyzed using far UV CD spectroscopy for their structure and EM for characterizing their morphology.

**Rheology of α-Syn gels**

To characterize the gels formed by WT and A53T α-Syn under LLPS conditions, both the proteins were incubated at 200 µM concentration in 20 mM sodium phosphate buffer (pH 7.4) in presence of 10% PEG. After the 30 days of incubation, the suspension transformed into amyloid hydrogel as confirmed by gel-inversion test. The rigidity of the WT and A53T α-Syn gels were compared by bulk rheology measurements. In brief, 400 µl of the hydrogel sample was analyzed for their stiffness using a parallel plate geometry (25 mm) rheometer (Anton Paar, Austria). The measurements were taken with 0.05% amplitude with a 0.5 mm gap size in dynamic oscillatory mode. Three decades (100-0.1) of angular frequency (ω) sweep was performed and the linear viscoelastic region was obtained from a preliminary strain sweep from 0.01% to 100%.

**FRAP data analysis**

After formation of rhodamine-labeled α-Syn liquid droplets under various LLPS inducing conditions (PEG, PD associated factors, phosphomimic S129 and familial A53T mutation), the slides containing the droplets were subjected to FRAP experiments at various time points. Briefly, appropriate region of interest (ROI) was chosen inside a single liquid droplet and the region was bleached with laser and further analyzed for its fluorescence recovery with time. Fluorescence recovery curves were constructed from the total intensity fluorescence values in the ROI for each frame corrected for the background and laser scanning bleaching. To calculate normalized fluorescence intensity, the following equation was used:

$$F(n) = \frac{[f(t)-f(b)]}{r} \quad (1)$$

Where, rate of photo-bleaching (r) = $f_c/f_{c0}$

$f_{c0}$ = fluorescence intensity of the ROI before photo-bleaching

$f_c$ = fluorescence intensity of the ROI after photo-bleaching

The mobile fraction (MF) of the protein molecule in the ROI is depicted by;

$$M.F = \frac{f\infty - f0}{fin - f0} \quad (2)$$

$f_\infty$= end value of fluorescence intensity post recovery

$f_{in}$= pre-bleach fluorescence intensity

$f_0$= post-bleach fluorescence intensity

The half-life of the recovery was calculated using curve fitting with exponential function with the help of software-generated algorithms in OriginPro 8.0 (OriginLab, USA).

To calculate the diffusion coefficient (D) from the half-life values, the following equation was used;

$$D = \frac{\omega^2}{4t_{1/2}} \quad (3)$$

Where, $\omega^2$ = area of the bleached region

And $t_{1/2}$ = half-life of recovery

### Time-resolved fluorescence spectroscopy analysis

The time-resolved fluorescence intensity decay curves were analyzed using methods described earlier (Swaminathan and Periasamy, 1996). The intensity decays were expressed as a sum of three exponentials as described below,

$$I(t) = \sum_{i=1}^{n} \alpha_i e^{-\left(\frac{t}{\tau_i}\right)} \quad (4)$$

I(t) stands for the intensity collected at magic angle (54.7°) at time t.

$\alpha_i$ stands for the amplitude of the fluorescence lifetime $\tau_i$ such that $\Sigma\alpha_i=1$. The mean fluorescence lifetime ($\tau_m$) was determined using the following equation,

$$\tau_m = \Sigma \frac{\alpha_i \tau_i}{\alpha_i} \quad (5)$$

The fluorescence anisotropy decay curves ($r(t)$) were analyzed using the equation,

$$r(t) = \frac{I_P(t) - G(\lambda) I_{pp}(t)}{I_P(t) + 2G(\lambda) I_{pp}(t)} \quad (6)$$

where $I_p$ stands for the fluorescence intensity when the emission polarizer was parallel to the excitation polarizer at time t.

$I_{pp}$ stands for the fluorescence intensity when the emission polarizer was perpendicular to the excitation polarizer at time t.

G($\lambda$) represents the geometry factor obtained for NATA at identical instrumental parameters as used for protein samples.

Analysis of the time resolved fluorescence anisotropy decay was performed by fitting $I_p$ and $I_{pp}$ as follows;

$$I_p(t) = I(t) \frac{\{1+2r(t)\}}{3} \quad (7)$$

$$I_{pp}(t) = I(t) \frac{\{1-r(t)\}}{3} \quad (8)$$

$$r\,(t) = r_0(\alpha_1 e^{-\frac{t}{\Phi 1}} + \alpha_2 e^{-\frac{t}{\Phi 2}}) \quad (9)$$

where $r_0$ is the fluorescence anisotropy when rotational diffusion is 0; $\alpha_i$ is the amplitude of the $i^{th}$ rotational correlation time (Φi) and;

$$\Sigma\, \alpha_i = 1$$

The $r_0$ value was previously reported to be 0.25 (Sahay et al., 2015) and was used for our calculations.

### Spatially-resolved fluorescence imaging-based FRET data analysis

Fluorescein maleimide (donor) and rhodamine maleimide C2 (acceptor) were separately labeled with Cys74 α-Syn according manufacturer’s protocol. 200 μM of fluorescein labeled α-Syn was mixed with equimolar rhodamine labeled α-Syn. 10% PEG was added to the mixture to initiate LLPS. After droplet formation, the fluorescence emission of acceptor was monitored upon excitation at the donor fluorescence excitation maxima. Rhodamine emission at 488 nm excitation from a single droplet containing both rhodamine and fluorescein labeled α-Syn is determined by dividing the emission spectrum of rhodamine at 532 nm excitation $(I^{A}_{\lambda_{ex}=532nm})$ with 5.4 factor as the extinction coefficient ($\epsilon^{rhodamine}_{532nm}$) of rhodamine. The emission from the tail of the fluorescein spectrum at the rhodamine peak ($I^{D,tail}_{\lambda_{ex}=488\ nm}$) has been added to get the precise emission from a droplet in absence of energy transfer. An intensity enhancement factor due to the energy transfer ($IEF_{ET}$) for a droplet containing both rhodamine and fluorescein has been defined from the emission of rhodamine at 488 nm excitation $(I^{A}_{\lambda_{ex}=488\ nm})$ in the following way:

$$IET_{ET} = \frac{I^{A}_{\lambda_{ex}=488nm}}{(0.185 \times I^{A}_{\lambda_{ex}=532\ nm} + I^{D,tail}_{\lambda_{ex}=488\ nm})} \qquad (10)$$

**Generation of stable HeLa cells expressing C4-α-Syn**

Plasmid construction and DNA manipulations were performed following standard protocols. The DNA sequences of all constructs were verified by sequencing prior to use. The C4-α-Syn-encoding plasmid (pcDNA5-C4-αSyn) was obtained by cloning the cDNA coding region of human α-Syn downstream the C4 tag (amino acids FLNCCPGCCMEP) into the destination vector pcDNA5-Dest/FRT/TO derived from pcDNA5/FRT/TO (Invitrogen) by two-step PCR. The open reading frame (ORF) constituted FLAG-StrepII-tetracysteine tag fused to the N-terminal of α-Syn gene.

The following primers were used to create pcDNA5-C4-αSyn;

<u>**FORWARD:**</u>

CCGGGACAAGTTTGTACAAAAAAGCAGGCTGCCACCATGGACTACAAGGACGACGA
CGACAAGTGGAGCCACCCCCAGTTCGAGAAGTTTCTCAATTGTTGTCCTGGCTGTTG
TATGGAACCTGATGTATTCATGAAAGGAC

<u>**REVERSE:**</u>

GGGGACCACTTTGTACAAGAAAGCTGGGTCTATTAGGCTTCAGGTTCGTAGTCTTGA
TACCCTTCCTCAGAAGGCATTTCATAAGCC

HeLaFlp-In/T-Rex cells (Invitrogen) were cultured in Dulbecco’s Modified Eagle Medium (DMEM, pH 7.3, GIBCO) supplemented with 10% fetal bovine serum (GIBCO), 4 mM L-glutamine, 100 U/ml penicillin and 100 mg/ml streptomycin. HeLa cell clones inducibly expressing C4-α-Syn were generated using the Flp-In/T-REx inducible expression system (Invitrogen) by co-transfecting the pcDNA5-C4-αSyn and pOG44 plasmids. Selection was carried out using 100 µg/ml hygromycin B. The medium was changed every two days until individual cell clones were established. For induction of C4-α-Syn expression, cells were treated with 20 ng/ml of doxycycline hyclate.

#### Live cell imaging

For live cell imaging experiments, HeLa cells were cultured as mentioned above in the confocal dishes (SPL Lifesciences, Korea). Post 16 h of culturing, 10 mM of ferric ammonium citrate was used to induce stress. For microtubule depolymerization, 16 µM of nocodazole was added to the cells and incubated for 2 h at 37 ºC in 5% $CO_2$. At given time-points, time-series images of cells stained with FlAsH–$EDT_2$ were captured using Zeiss Axio-Observer Z1 laser scanning confocal microscope (inverted) with 63X 1.4 NA oil immersion objective. FlAsH was excited with an Ar-ion laser at 488 nm, and detected with a 515 long-pass filter. Image processing and quantification was performed with Imaris 7.6.4 (BitPlane, Switzerland) and ImageJ (NIH).

#### Single Particle tracking

Single droplets inside the cell were identified and tracked as spots using Imaris 7.6.4 software (BitPlane, Switzerland). Using the autoregressive algorithm, the particle trajectory was generated. The track straightness, time-dependent positions and velocities for each droplet were determined. Mean squared displacement (MSD) analysis was done using *msdanalyzer* package (Tarantino et al., 2014) in MATLAB. We computed mean squared displacement ($r^2$) as a function of time (t); and from the statistical physics of particle movement, we expect power-law motion with $r^2$ proportional to t^ α, where diffusion exponent (α) is a parameter that can be obtained from the experimental data. Using the values of α, we characterized different types of motion. MSD curve for the trajectories were fitted to the quality of $R^2>0.8$ for generating the values of α. α=1 signifies free diffusive behavior of the particle, a value closer to 2 signifies active transport, while a value <1 refers to confined motion. The straightness of the path generated for each particle is defined as the ratio between displacement and length of track traversed by the particle. The value closer to 1 represents a more directed motion of the droplet. The track straightness and average velocity of each particle over time was calculated and plotted as a histogram using MATLAB 2018b.

#### Immunoprecipitation

3 x $10^6$ HeLa C4-α-Syn cells were used to perform immunoprecipitation. Untreated and 10 mM ammonium ferric citrate treated cells were collected at 24 h and 48 h. The cells were lysed in RIPA buffer with constant agitation at 4 °C for 30 min, and later centrifuged at 12,000 rpm for 20 min. Supernatant was quantified using the Bradford assay. Equal quantities of proteins were incubated with 2 µg of anti-FLAG antibody (Sigma) overnight at 4 °C on a rotating shaker. 50 µl Protein G sepharose beads (catalog no. P3296, Sigma, USA) were incubated with antibody-lysate mixture for 3 h at 4 °C to pull down the immunoprecipitated (IP) fraction. The IP fraction was eluted with 100 µl of 0.2 M glycine buffer, pH 2.6, and was then neutralized by equal volume of 50 mM Tris–HCl, pH 8.0.

**Dot blot**

4 μl of IP sample was spotted onto a polyvinylidene difluoride (PVDF) membrane (Millipore) and allowed to air dry. The membrane was fixed with 0.4% paraformaldehyde for 45 minutes. The membrane was treated with 5% non-fat skimmed milk powder in Tris-buffered saline with 0.1% Tween-20 (TBST) for 1 h at room temperature. The blots were rinsed with TBST and incubated with primary antibody (anti-FLAG, 1:1000 or OC, 1:2000) for overnight at 4 °C on a rocker. After incubation, the blots were washed three times with TBST for 10 min each, followed by 1 h incubation with horseradish peroxidase conjugated secondary antibody, 1:6000 (Calbiochem, USA) for 1 h at room temperature. Membranes were washed thrice with TBST, each for 10 min and signals were detected with SuperSignal West Femto kit (Thermo Scientific).

**Immunofluorescence**

FlAsH stained cells were washed with PBS (phosphate-buffered saline), pH 7.4 and fixed using 4% paraformaldehyde for 10 min. The fixed cells were permeabilized using 0.3% Triton X-100 in PBS for 10 min, and further nonspecific antigen sites were blocked using 2% BSA in PBS with 0.2% Tween-20 (PBST). The coverslips were incubated with primary antibody for α-tubulin (1:500) overnight at 4 °C in a humidified chamber and later washed three times with PBST. The coverslips were further incubated with goat anti-rabbit Alexa Fluor-555 (1:1500) secondary antibody for 2 h at room temperature in a humidified chamber. The coverslips were washed with PBST and the nucleus was counter stained with 1 μg/ml DAPI. The coverslips were mounted using 1% DABCO (1,4-diazabicyclo-[2.2.2]octane, Sigma) in 90% glycerol. Imaging was done with Axio-Observer Z1 laser scanning confocal microscope (Zeiss) with iPlan-Apochromat 63X/1.4 NA oil immersion objectives and processed with ImageJ (NIH).

**ProteoStat binding assay and flow cytometry analysis**

For imaging experiments, treated and untreated cells were fixed with 4% paraformaldehyde and permeabilized with 0.2% Triton X-100. In 1 ml of assay buffer (provided in the kit), 0.5 μl of ProteoStat (5 μM, Enzo Life Sciences, NY, USA) was added. 5 μM MG-132 treated cells treated were used as positive control. The cells were stained and imaged using confocal microscopy. For flow cytometric analysis, $\sim 1 \times 10^6$ cells were stained using ProteoStat, and analyzed using a BD FACS Aria flow cytometer using Texas Red channel (BD Biosciences, CA, USA). All of the experiments were performed at least three times. Flow cytometry data was analyzed by comparison of mean fluorescence, to compute aggresome propensity factor (APF), given as $APF = 100 \times ((MFI_{treated} - MFI_{untreated})/MFI_{treated})$, where $MFI_{treated}$ and $MFI_{control}$ are the mean fluorescence intensity (MFI) values from untreated and iron-treated samples. Aggresome propensity factor (APF), a unit-less term, provides a measure for dye accumulation within the cells in presence of proteasome inhibitor or inducer of protein aggregation.

**Statistical analysis**

The statistical significance was calculated by one-way variance analysis and subsequently, by the Student−Newman−Keuls multiple-comparison post hoc test (*$p \leq 0.05$; **$p \leq 0.005$;***$p \leq 0.001$; NS $p > 0.05$, NS: non-significant).

### RESULTS

#### Site-specific conformational changes and dynamics during LLPS

To gain insight into the domain involvement of α-Syn in LLPS at the residue level, two-dimensional [$^{15}N$-$^{1}H$] HSQC spectra were recorded for four days at an interval of 24 h and 20$^{th}$ day for WT and A53T α-Syn. The narrow signal dispersion in the direct dimension ($H^N$) indicated the disordered state of the proteins even in the presence of 10 % PEG (Figure S5). Upon incubation at 37 °C, WT α-Syn showed a gradual decrease in the intensities in the residues of the N-terminus (V2-A27, V37-K43, H50-E57) and non-amyloid-β component (NAC) regions (residues V74-V82 and A89-K97) over initial four days (Figures S5A and C). In case of A53T, immediately after 24 h, residues V3-K34 exhibited a drastic decrease in the intensities as compared to the other residues of N-terminus (36-58), NAC region (T75-G84 and A89-K97) and C-terminus (G111-L113 and D119-N122) (Figures S5B and D). After 48 h, amide cross peaks of V3-G14 disappeared completely due to line broadening; whereas a drastic decrease was observed for A89-K97, which completely disappeared after 96 h (Figure S5B) highlighting a critical role of these N-terminal and NAC domain residues in LLPS. On 20$^{th}$ day, for both WT and A53T, these peaks completely disappeared and new peaks were visible, highlighting a distinct conformation of α-Syn. The comparison of the intensity profile of WT and A53T residues after 48 h showed the residues in the N-termini of A53T possess a drastic decrease in intensity compared to WT during LLPS (Figures S5B and D). This data suggests that these N-terminal residues probably are important for promoting rapid phase separation of A53T mutant compared to WT α-Syn.

Although the intermolecular interaction of the protein during the phase separation is clear from the solution state ensemble FRET measurements, it is difficult to delineate the characteristics of domain interaction at a single droplet level. Therefore, using spatially-resolved fluorescence imaging and spectroscopy, we determined intermolecular FRET in a single liquid droplet. The proteins were individually labeled with fluorescein-5-maleimide and rhodamine red C2 maleimide at 74$^{th}$ position of cysteine, and equimolar concentration of these was mixed in 20 mM phosphate buffer (pH 7.4) in the presence of 10% PEG. The emission from the droplets was collected in two energetically separate detection channels (515−565 nm and 590−700 nm) to selectively detect fluorescein (green, false color) and rhodamine (red, false color). The quantitative energy-mapped fluorescence image shows the presence of both the fluorescein and the rhodamine-labeled protein in the droplets. However, there was significantly increased emission in the rhodamine detection channel compared to the fluorescein channel highlighting the possibility of energy transfer between the FRET pair or their preferred relative amounts in the droplets. To explore this, the emission spectrum of spatially separated individual droplets containing both the fluorescein and rhodamine was recorded under dual-wavelength excitation. First, the presence of rhodamine (acceptor) was detected at 532 nm excitation followed by the 488 nm excitation for the fluorescein (donor). The emission at 488 nm excitation from a droplet (Figure 5H, orange line) showed two distinct maxima due to the fluorescein ($I_{max}^{em} \approx 520nm$) and rhodamine ($I_{max}^{em} \approx 583nm$). It should be noted that the absorption coefficient ($\epsilon$) of rhodamine at 532 nm is ~5.5 times greater than at 488 nm ($0.185 \times \epsilon_{532nm}^{rhodamine} \approx \epsilon_{488nm}^{rhodamine}$) and fluorescein can barely absorb at rhodamine excitation (532 nm). To quantify energy transfer, we introduced a parameter defined as the intensity enhancement factor due to the energy transfer ($IEF_{ET}$, refer to Supplementary Methods), and calculated it for five individual liquid droplets over three different days to probe the change of apparent closeness. The $IEF_{ET}$ spectrum shows that energy transfer increases with the days (Figure S6C), suggesting a decrease in the apparent distance of the 74$^{th}$ position in the liquid droplet. In addition, the increased broadening of the spectrum at the 24$^{th}$ day suggests the formation of protein aggregates, which corroborates with amyloid aggregate formation in the droplet after ~20 days.

### FIGURE LEGENDS

**Figure S1: α-Syn undergoes liquid-liquid phase separation (LLPS) *in vitro***

**A**. DIC images of phase separated droplets in the presence of various concentrations of protein and PEG showing the regime of α-Syn LLPS. Higher concentration of either the protein or the molecular crowder (PEG-8000) or both could induce liquid phase separation of α-Syn. **B**. Light scattering measurements to quantify LLPS of α-Syn. Scattering values at 350 nm (triplicate measurements at each time-point) showing increased scattering, post 48 h, for 200 μM α-Syn incubated with 10% PEG-8000. Only α-Syn or PEG did not show any considerable light scattering. **C**. Liquid droplet formation by α-Syn using FITC-labeled α-Syn. The orthogonal Z-sectioning of the FITC-labeled α-Syn droplets confirms the spherical nature, and presence of α-Syn molecules inside the droplets. **D**. α-Syn LLPS regime at two different pHs (7.4 and 5.4). DIC images of the α-Syn phase separated droplet formed at pH 7.4 and 5.4 at different protein concentrations. At pH 5.4 (close to the protein's pI (4.7)), the critical concentration of α-Syn LLPS reduces to 5 μM in the presence of 10% PEG and 10 μM in absence of the crowding agent. **E and F**. α-Syn LLPS carried at different temperature and the reversibility of the droplets. (E). DIC images showing temperature-dependent LLPS of α-Syn at 4, 18, 25 and 37 °C. (F). Top panel: DIC images demonstrating the time-dependent disappearance and subsequent reappearance of α-Syn droplets when the droplets (formed after 2 days incubation) were first heated up to 65°C and then allowed to cool down to 37°C, respectively. The bottom panel represents 20 days old droplets that, on contrary, are resistant to change in temperature and is non-reversible. No change was observed when the temperature was shifted to 4 °C.

**Figure S2**: **PD-associated conditions promote α-Syn LLPS**

**A.** TEM images of α-Syn fibrils formed in the presence of 50 μM $Cu^{+2}$, 50 μM $Fe^{+3}$, 1 mM liposome with 200 μM protein. **B.** DIC images of 200 μM α-Syn showing liquid droplet formation in presence of different concentrations (mentioned in the figure) of the additives. It is important to note that in presence of 100 μM $Cu^{+2}$, $Fe^{+3}$ and >1 mM liposome, α-Syn showed aggregated structures under the microscope. Right panel shows LLPS at 5 μM protein concentration in the presence of indicated additives. **C.** DIC images of α-Syn (200 μM, 10% PEG) in presence of 200 μM dopamine showing \no phase separation even after 7 days of incubation at 37°C. **D**. Light scattering measurements after 48 h for 200 μM protein in 10% PEG-8000 showing higher increase in the light scattering for A53T compared to WT protein. **E.** Quantification of the number and size of liquid droplets at different time points for WT and A53T α-Syn. Number of droplets (n=200) were counted from five different microscopic fields (63X magnification), values represent mean ± s.e.m. **F**. The % fluorescence recovery of liquid droplets formed at day 2 in presence of additives after photobleaching (FRAP) analysis.

**Figure S3: Isolation and analysis of monomer, oligomer and fibril fractions formed during LLPS**

**A.** Schematic showing isolation procedure for soluble (LMW), oligomeric and fibrillar α-Syn during LLPS. **B.** Isolated LMW α-Syn from the LLPS system in presence of 10 % PEG showing random coil structure (~198 minima) for all the time points. Nonetheless, the extent of the signal decreased with time due to changes in α-Syn monomer over time. Oligomer isolated at day 5 (d5) showed random coil whereas from d10 to d30 showed α-helix (two minima at ~208 nm and at 222). For fibril component, α-Syn showed mostly β-sheet rich structure (~218 nm) at all the time points of isolation. **C.** The morphology using TEM analysis of LMW, oligomer and fibril isolated at various time points showing amorphous, globular and fibrillar-aggregate, respectively. 50 μM of protein concentration was used for TEM.

**Figure S4: Effect of mutations on LLPS and aggregation of α-Syn**

**A**. Aggregation kinetics for WT and A53T under LLPS conditions (200 μM α-Syn+10% PEG, no agitation, incubated at 37°C) monitored by ThT assay. **B**. Representative images of A53T α-Syn droplets showing positive ThioS binding after day 2 of incubation and droplet maturation. TEM image reveals the presence of droplets at the early days of incubation and formation of amyloid fibrils at the late stage (day 30). **C.** The gel inversion test for A53T protein solution in presence of 10% PEG after 30 days of incubation depicting hydrogel formation (left) and the corresponding scanning electron microscope (SEM) showing hydrogel networks (right). The inset showing bulk rheology of WT and A53T α-Syn hydrogel with higher storage modulus (G') than loss modulus for both the proteins; confirming the gel-state. **D.** Aggregation kinetics under rotating conditions for 200 μM of α, β, γ and core-Syn monitored using ThT fluorescence. β- and γ-Syn did not show any aggregation during the indicated incubation time. **E**. LLPS behavior of β, γ and core-Syn (200 μM protein, 10% PEG, pH 7.4, 37°C, no agitation) showing no LLPS for β or γ-Syn. The core α-Syn (30-110) shows faster LLPS than full length WT α-Syn (which takes 2 days in identical conditions). At day 6, core α-Syn (30-110) started to form aggregate-like structures (white arrow). **F**. TEM images of α, β, γ and core-Syn after one month of incubation under LLPS condition showing fibril formation for full-length and core α-Syn. However, β or γ-Syn did not show any amyloid formation.

**Figure S5: Domain involvement of α-Syn during LLPS monitored by NMR spectroscopy**
**A, B.** Two-Dimensional [$^{15}N$-$^{1}H$] HSQC spectra (red) of α-Syn at the beginning of incubation LLPS (day 0) overlapped with spectra obtained after day 1 (blue), day 2 (green), day 3 (sky blue) day 4 (teal) and day 20 (magenta) for WT (A) and A53T mutant proteins (B). **C, D**. Intensity ($I/I_0$) profile of amide cross-peaks from [$^{15}N$-$^{1}H$] HSQC spectra of α-Syn on day 1 (red), day 2 (cyan), day 3 (blue) and day 4 (green) normalized against intensity of peaks on day 0 (grey) for WT (C) and A53T mutant (D). Insets show DIC microscopic images for liquid droplets formed in the sample for NMR study providing direct comparison. The data show drastic decrease in the NMR signal intensities for the residues at the N-terminus and NAC domain during LLPS, and further maturation of liquid droplets. NMR signal decays at much more extent for A53T compared to WT protein showing faster liquid droplet formation by A53T.

**Figure S6**: **Spectroscopic analysis for monitoring domain involvement during α-Syn LLPS**
**A.** Time-resolved fluorescence intensity and anisotropy decay by four α-Syn Trp mutants at different positions (residue 3W, 71W, 124W and 140W). **B.** Fluorescence spectroscopy based intermolecular FRET spectra of single Trp at 3$^{rd}$, 71$^{st}$ and 124$^{th}$ position of α-Syn as donors and single Cys-DTNB at 3$^{rd}$, 74$^{th}$ and 124$^{th}$ position of α-Syn as acceptors. The Trp fluorescence emission decreased rapidly with time for all the positions, which suggest positive fluorescence energy transfer from Trp to DTNB. However, the extent of FRET signature was higher for 3W-3C (DTNB) and the 71W-74C (DTNB) compared to 124W-124C (DTNB) in early days of LLPS. The fluorescence intensities were normalized with the Trp only controls without any FRET. **C.** Spatially-resolved fluorescence spectra of single phase separated droplets at indicated time points at the 74$^{th}$ position. The solid lines represent rhodamine emission when excited at 488 nm for five individual droplets; whereas dotted lines represent no FRET for those of the five respective droplets. The thick line is the average of these five spectra when FRET occurs. These spectra were utilized to extract $IEF_{ET}$ on different days. The spectral broadening at day 24 (d24) suggests the formation of amyloid aggregates.

**Figure S7. *In cell* analysis of liquid-like condensates of α-Syn**
**A**. Fluorescence and DIC merged images of iron-treated HeLa cells expressing C4-α-Syn stained with FlAsH-$EDT_2$. The images were captured at 24 h and 48 h post treatment. The white dotted segmentation marks the nuclear boundary. **B**. Quantification of the droplet number in the iron-treated cells. Number of cells for droplet count from three independent experiments are n = 20.

Data is presented as mean ± s.e.m., ***$p \leq 0.001$. **C**. α-Syn liquid droplets do not associate with membranes or lipid. Representative confocal images of Nile red, MitoTracker and LysoTracker-red dye staining of cells showing that the FlAsH stained C4-α-Syn droplets are not associated with the cellular lipid droplets or membrane-bound organelles, mitochondria and lysosomes. The images were acquired at 63X magnification. Scale bar: 10 μm. **D**. Mean squared displacement (MSD) curve for the droplet particles at 24 h (blue) and 48 h (red) of iron-treatment. The curve for 24 h shows a slightly positive curvature, while at 48 h, the curve is mostly linear. The plot is generated with MATLAB-based single particle tracking analysis of the time-series confocal images (n = 5) processed using IMARIS version 7.6.4. **E**. Representative trajectories for droplets at 24 h with $\alpha < 1$, $\alpha = 1$ and $\alpha > 1$. **F**. Straightness of the trajectories traversed by the droplets showing a shift in the population towards less straightness for α-Syn droplets at 48 h and with nocodazole treatment at 24 h. **G**. The correlation plots of straightness of the particles and their corresponding trajectories (α). **H**. A plot for distribution of $V_{rms}$ (root mean square of the velocity) of the particles at three conditions showing a reduction in $V_{rms}$ for 48 h droplets compared to 24 h droplets. Further, nocodazole treatment of cells also reduces the $V_{rms}$ of droplets formed at 24 h.

**Figure S8. α-Syn liquid-liquid phase separation and liquid-to-solid transition involves aggregation and aggresome formation *in cells***

**A**. Immunoprecipitation of α-Syn using anti-FLAG antibody from cell extracts of untreated and treated (with $Fe^{3+}$) cells at given time points. The dot blots were probed with amyloid-specific OC antibody. Anti-FLAG probing was done to ensure equal loading of the immunoprecipitates. **B**. Percentage cell viability measured using FITC-Annexin V - PI staining and FACS analysis for α-Syn expressing and non-expressing HeLa cells those untreated and treated with the $Fe^{3+}$stressor. No significant difference in the percentage cell viability is observed when compared among the two groups of cells. **C**. Aggresome detection using ProteoStat-dye staining. Confocal images of the iron-treated and untreated cells, at various time points, stained with FlAsH-$EDT_2$ (α-Syn) and ProteoStat dye. DAPI (blue) is for nuclear staining. **D**. Aggresome propensity factor (measured by ProteoStat) determined using FACS analysis (refer to supplementary method). Proteasomal inhibitor, MG132, was used as a positive control for aggresome induction. Aggresome propensity factor considerably increases at 48 h compared to earlier time-points of iron treatment.

### Supplementary video legends

#### Video S1- Liquid-Liquid phase separation of α-synuclein *in vitro*

Time-lapse movie showing formation of α-synuclein liquid droplets *in vitro*. The droplet size increases over time due to fusion and Ostwald ripening.

#### Video S2- α-Synuclein forms liquid droplets in cells.

A time-lapse movie showing formation of α-synuclein droplets in cells under oxidative stress. The droplets are highly dynamic in nature and fuse together to form larger droplets indicating their liquid-like behavior.

#### Video S3- α-Synuclein droplets cluster at the perinuclear region upon maturation.

A time-lapse movie of α-synuclein droplets after 48 hours of oxidative stress in HeLa cells showing their perinuclear localization. These droplets possess restricted movement compared to droplets formed at earlier time point, indicating their maturation and transformation into to a solid-like state.

#### Video S4- Disruption of microtubules leading to clumping of α-synuclein droplets in cell.

Time-lapse movie of α-synuclein droplets after nocodazole treatment in HeLa cells showing clumping and restricted movement of droplets compared to nocodazole untreated cells. The data indicating microtubules assist α-Syn droplets movement in cells.

### REFERENCES


Kumar, R., Das, S., Mohite, G.M., Rout, S.K., Halder, S., Jha, N.N., Ray, S., Mehra, S., Agarwal, V., and Maji, S.K. (2018). Cytotoxic Oligomers and Fibrils Trapped in a Gel-like State of alpha-Synuclein Assemblies. Angew. Chem. *57*, 5262-5266.

Sahay, S., Ghosh, D., Dwivedi, S., Anoop, A., Mohite, G.M., Kombrabail, M., Krishnamoorthy, G., and Maji, S.K. (2015). Familial Parkinson disease-associated mutations alter the site-specific microenvironment and dynamics of alpha-synuclein. J.Biol.Chem. *290*, 7804-7822.

Singh, P.K., Kotia, V., Ghosh, D., Mohite, G.M., Kumar, A., and Maji, S.K. (2013). Curcumin modulates alpha-synuclein aggregation and toxicity. ACS Chem.Neurosci. *4*, 393-407.

Tarantino, N., Tinevez, J.Y., Crowell, E.F., Boisson, B., Henriques, R., Mhlanga, M., Agou, F., Israel, A., and Laplantine, E. (2014). TNF and IL-1 exhibit distinct ubiquitin requirements for inducing NEMO-IKK supramolecular structures. J. Cell Biol. *204*, 231-245.

Volles, M.J. and Lansbury, P.T. (2007) Relationships between the sequence of alpha-synuclein and its membrane affinity, fibrillization propensity, and yeast toxicity. J. Mol. Biol. 366, 1510-1522.

Figure S1

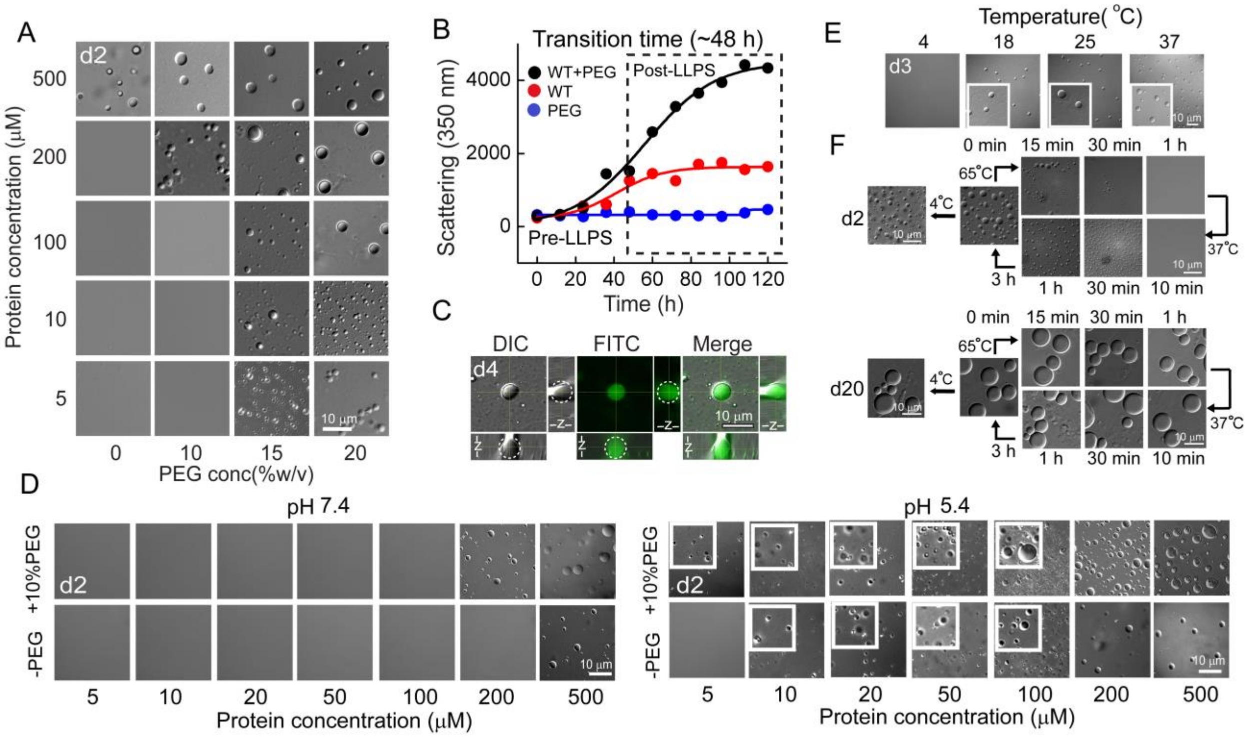
A
d2
500
200
100
10
5
Protein concentration (μM)
0
10
15
20
PEG conc(%w/v)
10 μm
B
Transition time (~48 h)
WT+PEG
WT
PEG
Post-LLPS
Pre-LLPS
Scattering (350 nm)
4000
2000
0
0
20
40
60
80
100
120
Time (h)
C
DIC
FITC
Merge
d4
10 μm
E
Temperature( °C)
4
18
25
37
d3
10 μm
F
0 min
15 min
30 min
1 h
65°C
4°C
d2
37°C
3 h
1 h
30 min
10 min
d20
D
pH 7.4
+10%PEG
-PEG
5
10
20
50
100
200
500
Protein concentration (μM)
pH 5.4
Protein concentration (μM)

Figure S2

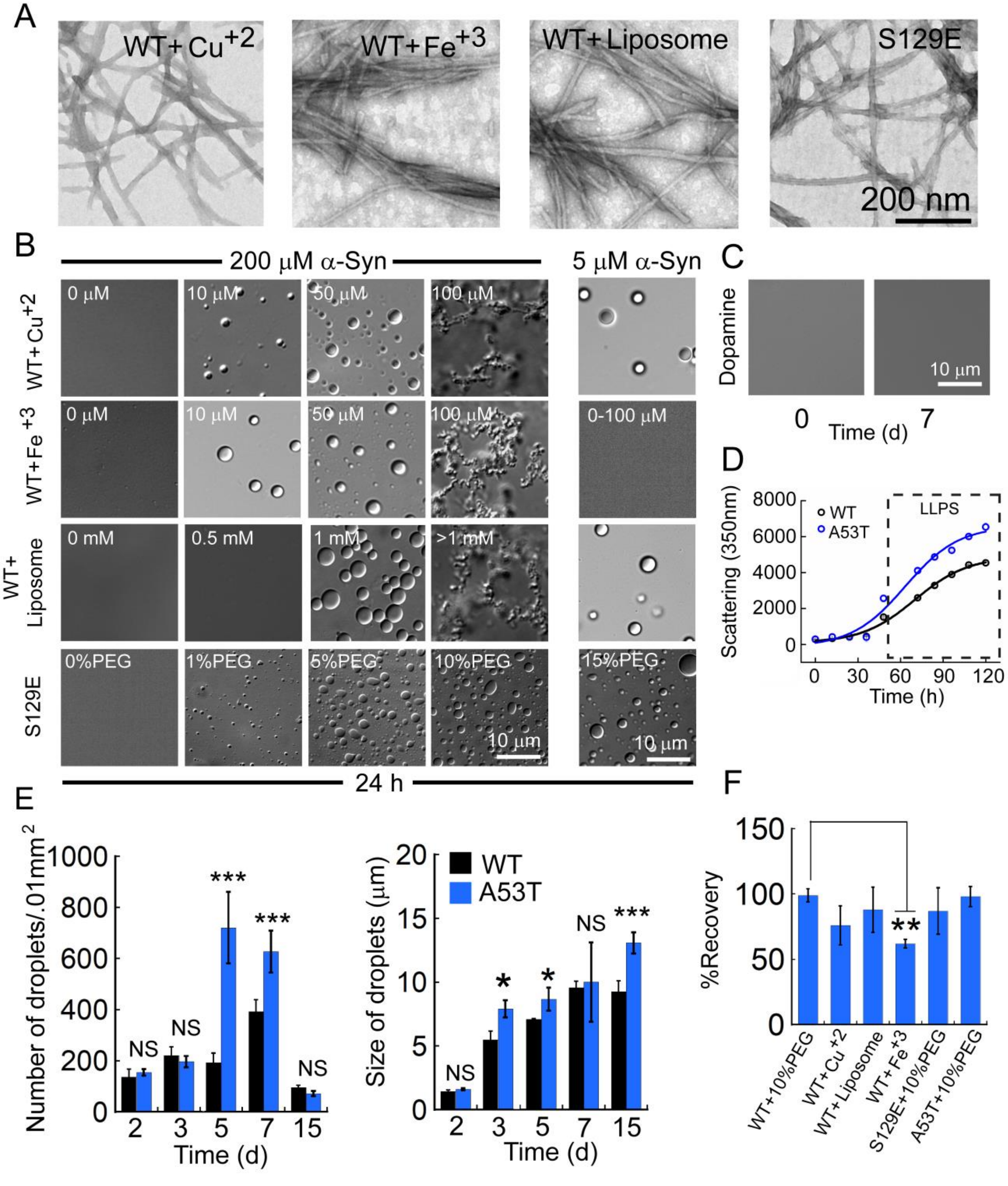
A
WT+Cu+2
WT+Fe+3
WT+Liposome
S129E
200 nm
B
200 μM α-Syn
5 μM α-Syn
WT+Cu+2
WT+Fe+3
WT+ Liposome
S129E
0 μM
10 μM
50 μM
100 μM
0-100 μM
0 mM
0.5 mM
1 mM
>1 mM
0%PEG
1%PEG
5%PEG
10%PEG
15%PEG
10 μm
24 h
C
Dopamine
10 μm
0
Time (d)
7
D
Scattering (350nm)
WT
A53T
LLPS
Time (h)
E
Number of droplets/.01mm2
Size of droplets (μm)
Time (d)
NS
***
*
F
%Recovery
**
WT+10%PEG
WT+Cu+2
WT+Liposome
WT+Fe+3
S129E+10%PEG
A53T+10%PEG

Figure S3

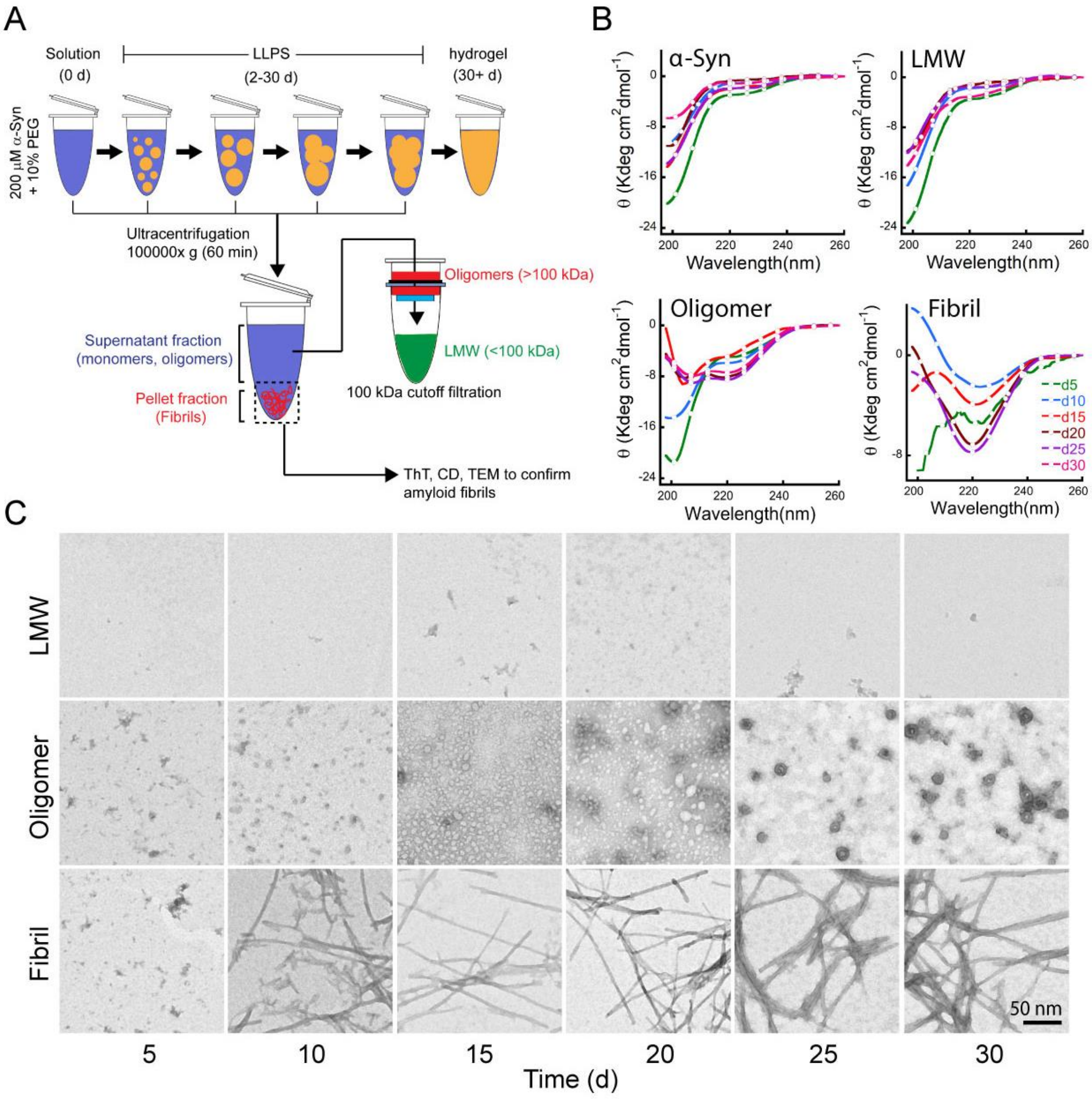
A
Solution
(0 d)
LLPS
(2-30 d)
hydrogel
(30+ d)
200 μM α-Syn
+ 10% PEG
Ultracentrifugation
100000x g (60 min)
Oligomers (>100 kDa)
LMW (<100 kDa)
Supernatant fraction
(monomers, oligomers)
Pellet fraction
(Fibrils)
100 kDa cutoff filtration
ThT, CD, TEM to confirm
amyloid fibrils
B
α-Syn
LMW
Oligomer
Fibril
θ (Kdeg cm²dmol⁻¹)
Wavelength(nm)
d5
d10
d15
d20
d25
d30
C
LMW
Oligomer
Fibril
50 nm
5
10
15
20
25
30
Time (d)

Figure S4

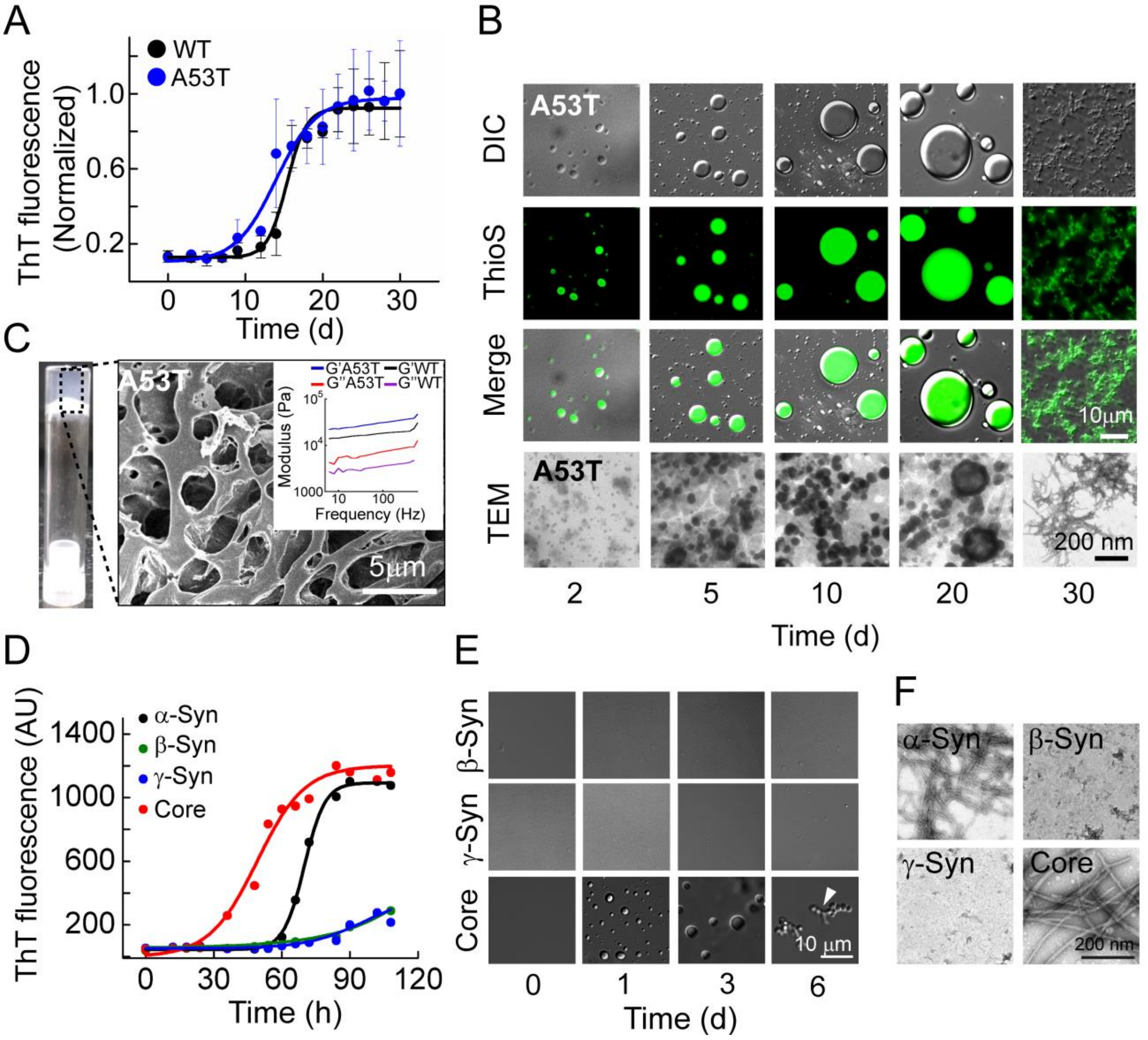
A
WT
A53T
ThT fluorescence (Normalized)
Time (d)
B
DIC
ThioS
Merge
TEM
A53T
10μm
200 nm
2
5
10
20
30
Time (d)
C
A53T
G'A53T
G'WT
G"A53T
G"WT
Modulus (Pa)
Frequency (Hz)
5μm
D
α-Syn
β-Syn
γ-Syn
Core
ThT fluorescence (AU)
Time (h)
E
β-Syn
γ-Syn
Core
10 μm
0
1
3
6
Time (d)
F
α-Syn
β-Syn
γ-Syn
Core
200 nm

Figure S5

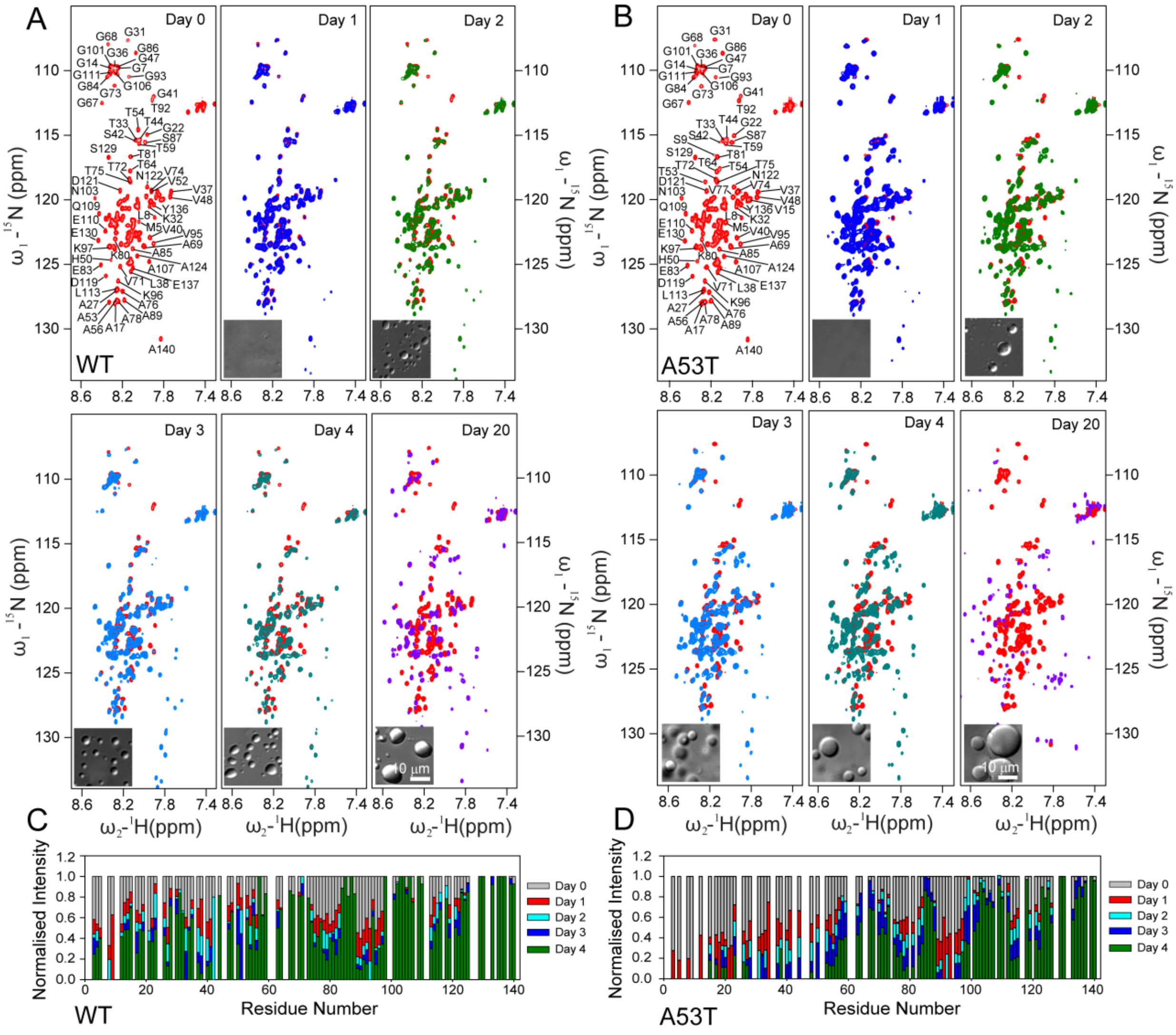

Figure S6

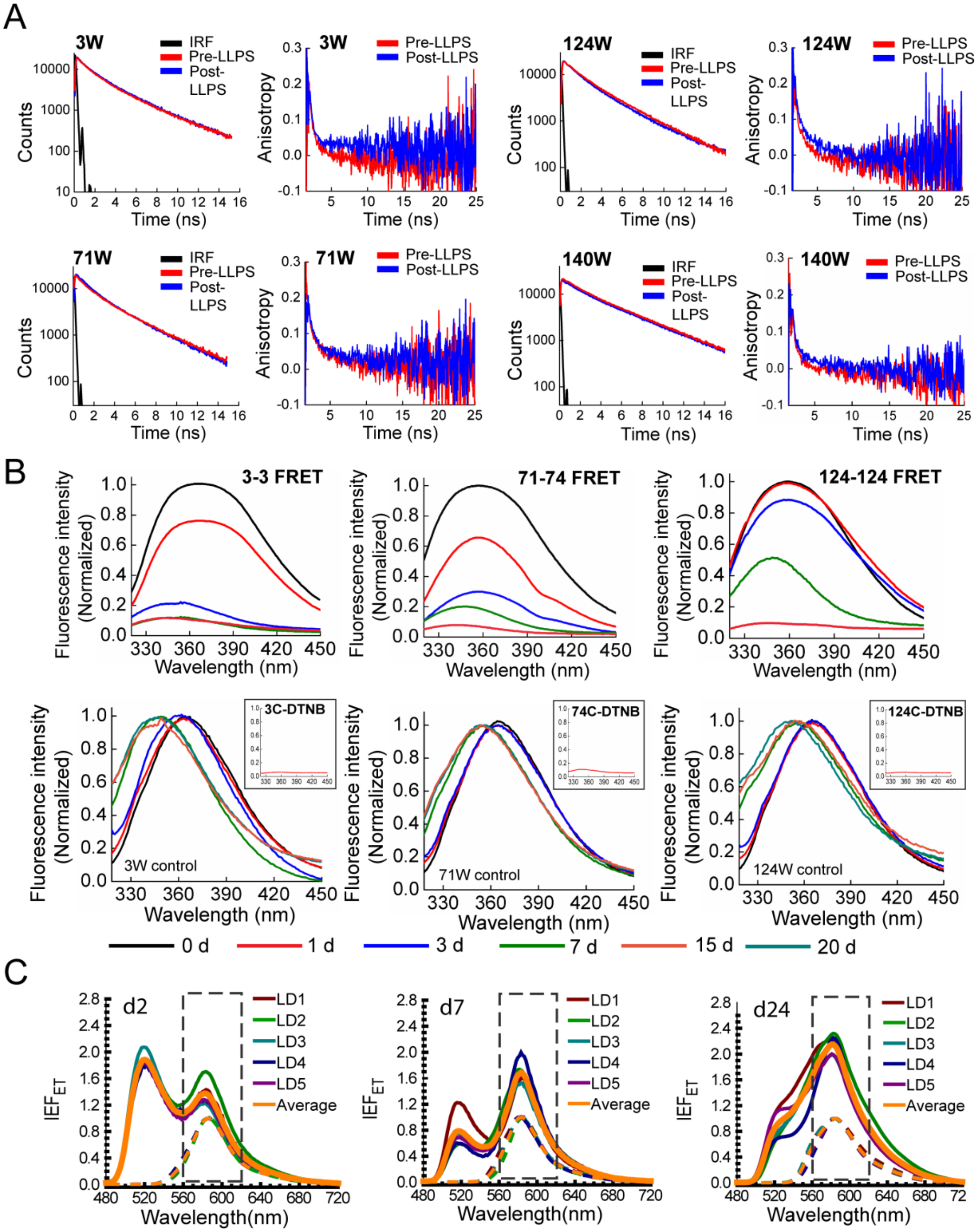
A
3W
IRF
Pre-LLPS
Post-LLPS
Counts
Time (ns)
Anisotropy
124W
71W
140W
B
3-3 FRET
71-74 FRET
124-124 FRET
Fluorescence intensity (Normalized)
Wavelength (nm)
3C-DTNB
74C-DTNB
124C-DTNB
3W control
71W control
124W control
0 d
1 d
3 d
7 d
15 d
20 d
C
d2
d7
d24
LD1
LD2
LD3
LD4
LD5
Average
$IEF_{ET}$
Wavelength(nm)

Figure S7

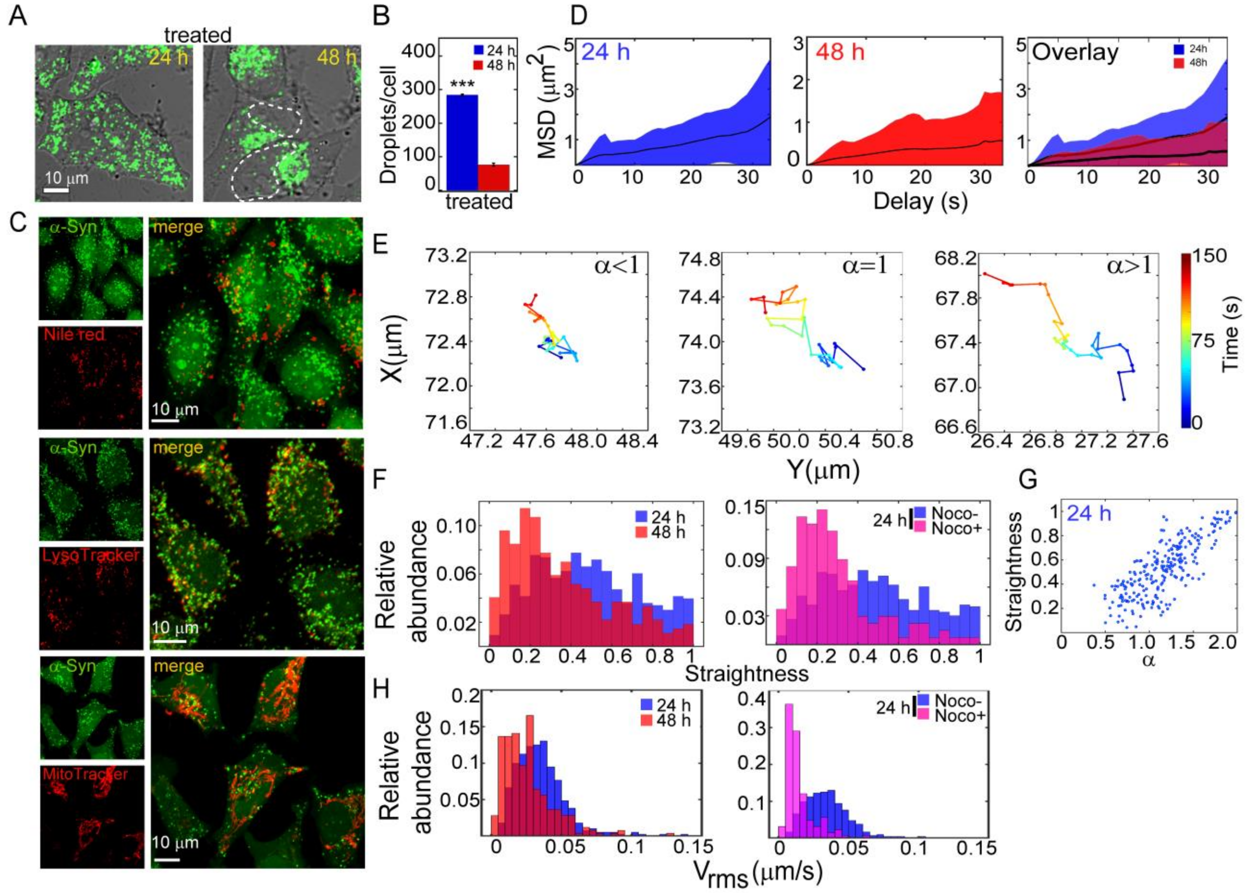

A
treated
24 h
48 h
10 μm
B
Droplets/cell
24 h
48 h
***
treated
C
α-Syn
merge
Nile red
LysoTracker
MitoTracker
10 μm
D
24 h
48 h
Overlay
MSD (μm2)
Delay (s)
E
α<1
α=1
α>1
X(μm)
Y(μm)
Time (s)
F
Relative abundance
Straightness
24 h
Noco-
Noco+
G
Straightness
α
H
Vrms (μm/s)

Figure S8

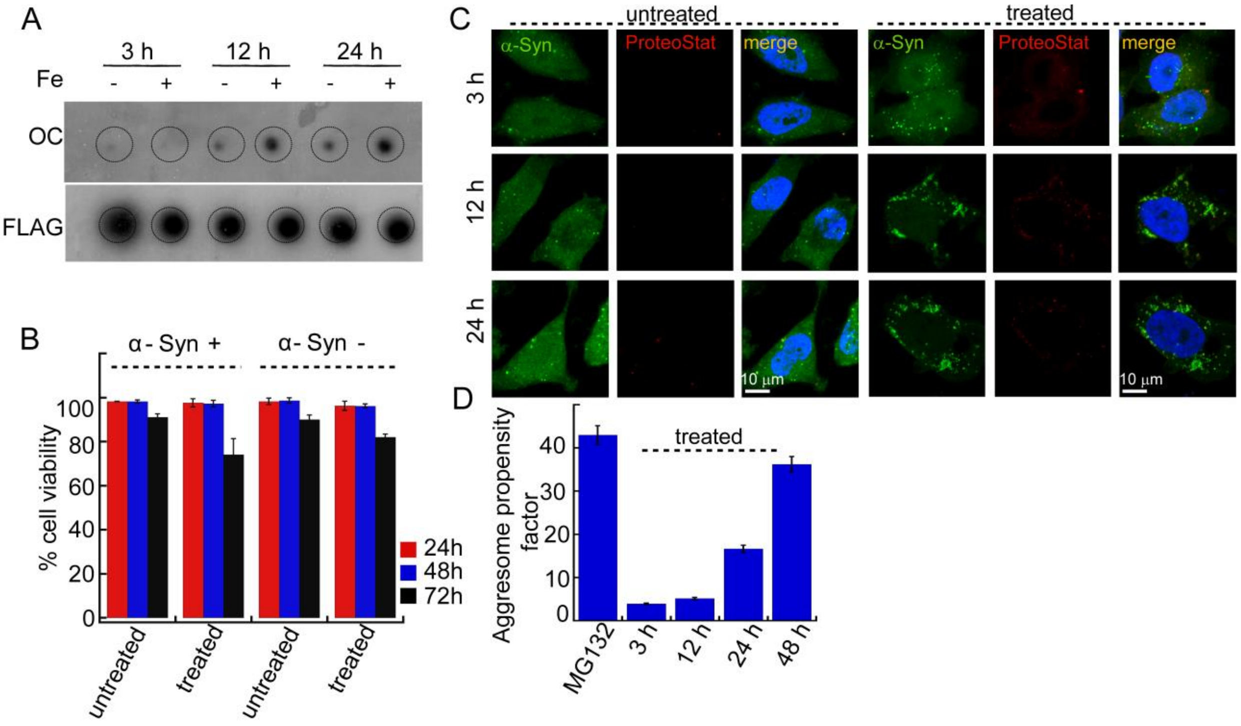

A
3 h
12 h
24 h
Fe
- + - + - +
OC
FLAG
B
α- Syn +
α- Syn -
% cell viability
100
80
60
40
20
0
24h
48h
72h
untreated
treated
untreated
treated
C
untreated
treated
α-Syn.
ProteoStat
merge
α-Syn
ProteoStat
merge
3 h
12 h
24 h
10 μm
10 μm
D
Aggresome propensity factor
treated
40
30
20
10
0
MG132
3 h
12 h
24 h
48 h
